# UGT76B1, a promiscuous hub of small molecule-based immune signaling, glucosylates N-hydroxypipecolic acid and controls basal pathogen defense

**DOI:** 10.1101/2020.07.12.199356

**Authors:** Sibylle Bauer, Dereje W. Mekonnen, Michael Hartmann, Robert Janowski, Birgit Lange, Birgit Geist, Jürgen Zeier, Anton R. Schäffner

## Abstract

Glucosylation modulates the biological activity of small molecules and frequently leads to their inactivation. The *Arabidopsis thaliana* glucosyltransferase UGT76B1 is involved in conjugating the stress hormone salicylic acid (SA) as well as isoleucic acid (ILA). Here, we show that UGT76B1 also glucosylates N-hydroxypipecolic acid (NHP), which is synthesized by FLAVIN-DEPENDENT MONOOXYGENASE 1 (FMO1) and activates systemic acquired resistance (SAR). Upon pathogen attack, Arabidopsis leaves accumulate two distinct NHP hexose conjugates, NHP-*O*-β-glucoside and NHP glucose ester, which are oppositely regulated by SA. *ugt76b1* mutants specifically fail to generate the NHP-*O*-β-glucoside, and recombinant UGT76B1 synthesizes NHP-*O*-β-glucoside *in vitro* in competition with SA and ILA. The loss of *UGT76B1* elevates the endogenous levels of NHP in addition to SA and ILA and establishes a SAR-like, primed immune status without pathogen infestation. The introgression of the *fmo1* background lacking NHP biosynthesis into *ugt76b1* abolishes the SAR-like resistance phenotype indicating an important function of UGT76B1-mediated NHP glucosylation in balancing the defense status. Our results further indicate that ILA promotes and SA finally executes the NHP-triggered immunity via the glucosyltransferase UGT76B1 as the common metabolic hub. Thus, UGT76B1 controls the levels of active NHP, SA, and ILA in concert to modulate plant immune signaling.

## INTRODUCTION

Defense against pathogens by plants is induced at the site of inoculation to combat the infection *in situ*, whereas uninfected plants lack induction or suppress inducible immunity. In addition, systemic acquired resistance (SAR) can be activated by systemic tissues protecting against subsequent infections (Vlot et al., 2009; Shah and Zeier, 2013; Fu and Dong, 2013; Klessig et al., 2018). A variety of small molecules are employed by plants to initiate and regulate these defense responses upon pathogen attack. Salicylic acid (SA) is a key signaling molecule required for both local and systemic defense. Upon pathogen infection, SA is synthesized primarily *via* the precursor isochorismate by ICS1/SID2 in chloroplasts and exported by EDS5 to the cytosol, where it initiates defense responses. A major route involves the establishment of a lower redox potential leading to NONEXPRESSOR OF PATHOGENESIS-RELATED GENES 1 (NPR1)-dependent changes in gene expression (Mou et al., 2003; Ding et al., 2018). Different SA conjugates accumulate in response to pathogen inoculation besides free SA, with glucoside formation being the dominant SA modification. SA glucosylation is considered crucial for controlling the level of unconjugated SA and thereby attenuating the defense responses (Vlot et al., 2009; Dempsey et al., 2011; Dempsey & Klessig, 2017). Three small-molecule glucosyltransferases of *Arabidopsis thaliana*, UGT74F1, UGT74F2 and UGT76B1, have been identified as SA glucosyltransferases (Dean and Delany, 2008; Song et al., 2008; von Saint Paul et al., 2011; Noutoshi et al., 2012; Li et al., 2015). Interestingly, UGT76B1 also glucosylates another small molecule, isoleucic acid (ILA; 2-hydroxy-3-methyl pentanoic acid) (von Saint Paul et al., 2011). ILA competitively inhibits SA glucosylation by UGT76B1. Therefore, the enhanced pathogen resistance synergistically induced by exogenous application of ILA and SA suggested the suppression of the attenuating SA glucosylation by ILA (Noutoshi et al., 2012; Bauer et al., 2020). The biosynthesis of ILA has not been resolved in plants; however, it is likely linked to the biosynthesis or catabolism of the branched-chain amino acid isoleucine via reduction of its deaminated derivative 2-keto-3-methyl pentanoic acid (Maksym et al., 2018).

The lysine-derived amino acid pipecolic acid (Pip) and its oxidized derivative N-hydroxypipecolic acid (NHP) are two other small molecules important to pathogen defense, in particular for the establishment of SAR (Návarová et al., 2012; Bernsdorff et al., 2016; Chen et al., 2018; Hartmann et al., 2018). Two major enzymes, AGD2-LIKE DEFENSE RESPONSE PROTEIN1 (ALD1), an L-Lys-α-aminotransferase, and SAR-DEFICIENT4 (SARD4), a reductase, are involved in Pip biosynthesis from lysine (Ding et al., 2016; Hartmann et al., 2017; Hartmann and Zeier, 2018). Characterization of Arabidopsis *ald1* mutants defective in the endogenous accumulation of Pip and exogenous feeding of Pip in various scenarios indicated its importance in plant defense responses and showed the Pip-dependency of SAR (Návarová et al., 2012; Vogel-Adghough et al., 2013). However, *fmo1* mutants were completely compromised in SAR and failed to acquire resistance in response to exogenous Pip feeding, suggesting the requirement of another molecule downstream of FLAVIN-DEPENDENT MONOOXYGENASE 1 (FMO1) for SAR (Mishina and Zeier, 2006; Návarová et al., 2012). FMO1 was then biochemically characterized *in vitro* and *in planta* as a pipecolate N-hydroxylase that catalyzes the conversion of Pip to NHP (Chen et al., 2018; Hartmann et al., 2018). Upon pathogen attack, NHP accumulates systemically in the Arabidopsis foliage. In addition, exogenous feeding with NHP potently activated SAR to bacterial and oomycete infection and abrogated the SAR-defect of the NHP-deficient *fmo1* mutant, indicating that NHP is the actual SAR-inducing compound of this resistance pathway (Chen et al., 2018; Hartmann et al., 2018). Similar to SA and ILA, also a glycosylated form of NHP was detected in Arabidopsis leaves after pathogen inoculation or exogenous application of NHP. However, the UGT involved in NHP hexoside formation and the functional implication of this conjugation remained elusive. It had been speculated that NHP hexoside could be an inactivated storage form and/or a mobile entity in SAR (Chen et al., 2018; Hartmann and Zeier, 2018).

Transcriptome analyses of *Arabidopsis* plants treated with exogenous Pip as well as of systemic SAR tissue revealed a strong induction of genes involved in SA, Pip and NHP biosynthesis (*e. g*. *ICS1*, *EDS5*, *ALD1*, *SARD4*, and *FMO1*). The strong upregulation of these genes indicates that the molecular implementation of the Pip-induced defense responses, including a positive feedback amplification of Pip, relies on links between these signaling molecules. Defense gene induction by Pip was dependent on functional *FMO1*, suggesting the requirement of NHP formation in SAR-related gene expression (Hartmann et al., 2018). The same Pip-, FMO1- and SAR-dependent expression pattern was also found for *UGT76B1*, the gene encoding the SA- and ILA-glucosyltransferase (Hartmann et al., 2018).

Therefore, we aimed at examining the role of UGT76B1 in the Pip-induced defense and its regulation *via* NHP. We addressed the effect of Pip application on *ugt76b1* loss-of-function mutants, which show enhanced resistance towards biotrophic pathogens, and on plants constitutively overexpressing UGT76B1, which are more susceptible than the wild type (von Saint Paul et al., 2011). A comparative metabolic analysis revealed that the wild type and UGT76B1-overexpressing plants accumulated an NHP-*O*-glucoside termed NHP-H2, which was completely absent from *ugt76b1*. This suggested a direct additional activity of UGT76B1 as an NHP glucosyltransferase, which could be confirmed *in vitro* and *in vivo*. Importantly, NHP-H2 was distinct from a previously reported NHP hexoside (Hartmann and Zeier, 2018), here termed NHP-H1, which is formed in a UGT76B1-independent manner. Mass spectrometry and digestion experiments suggest that this conjugate represents a NHP glucose ester. We show that UGT76B1 orchestrates the glucosylation of the three defense activators SA, ILA, and NHP and thereby controls the basal defense. The loss of UGT76B1 leads to a SAR-like, NHP-dependent enhanced immune status.

## RESULTS

### The formation of an NHP hexoside is abolished by *ugt76b1*

Previous studies have shown that exogenous application of pipecolic acid acid (Pip) strongly enhances the transcripts of the Arabidopsis SA glucosyltransferase encoding *UGT76B*1 along with several SAR-related genes including genes involved in SA and NHP biosynthesis. This transcriptional response to Pip was lost by *fmo1* plants indicating a positive impact of FMO1 or its product NHP on *UGT76B1* expression (Hartmann et al., 2018; Hartmann and Zeier, 2019). To conversely investigate a potential impact of UGT76B1 on the metabolic response to exogenous Pip, we performed comparative LC-MS-based metabolite analyses of leaf extracts of a *ugt76b1* loss-of-function mutant, a UGT76B1-overexpressing line, and the wild type. Exogenous Pip application was associated with a considerable uptake and relocation of Pip to rosette leaves by all three genotypes (Figure 1A). In addition, wild-type and *UGT76B1*-overexpressing (OE) plants accumulated low amounts of the Pip derivative NHP in response to Pip application (Figure 1B). Notably, the leaves of *ugt76b1* plants showed a by far higher NHP level upon Pip treatment than the other lines. Moreover, untreated *ugt76b1* exhibited a strong endogenous accumulation of both Pip and, in particular, of NHP (Figure 1A and 1B). This indicated a role of UGT76B1 in suppressing the levels of both Pip and NHP. Furthermore, a metabolite with *m/z* 308.1346 was enhanced upon Pip application by the wild type and UGT76B1-overexpressing plants (Figure 1C). The LC-MS-derived mass spectrum with a [M+H]^+^ ion at *m/z* 308.1346 (± 0.005) and a main fragment ion at *m/z* 146.0814, which is characteristic for a [NHP+H]^+^ ion, supported an NHP hexoside structure for this substance (Figure 1G; Chen et al., 2018). Intriguingly, this putative hexoside was undetectable in leaf extracts of the *ugt76b1* mutant, strongly suggesting an indispensable role of UGT76B1 in its formation (Figure 1C).

**Figure 1.**
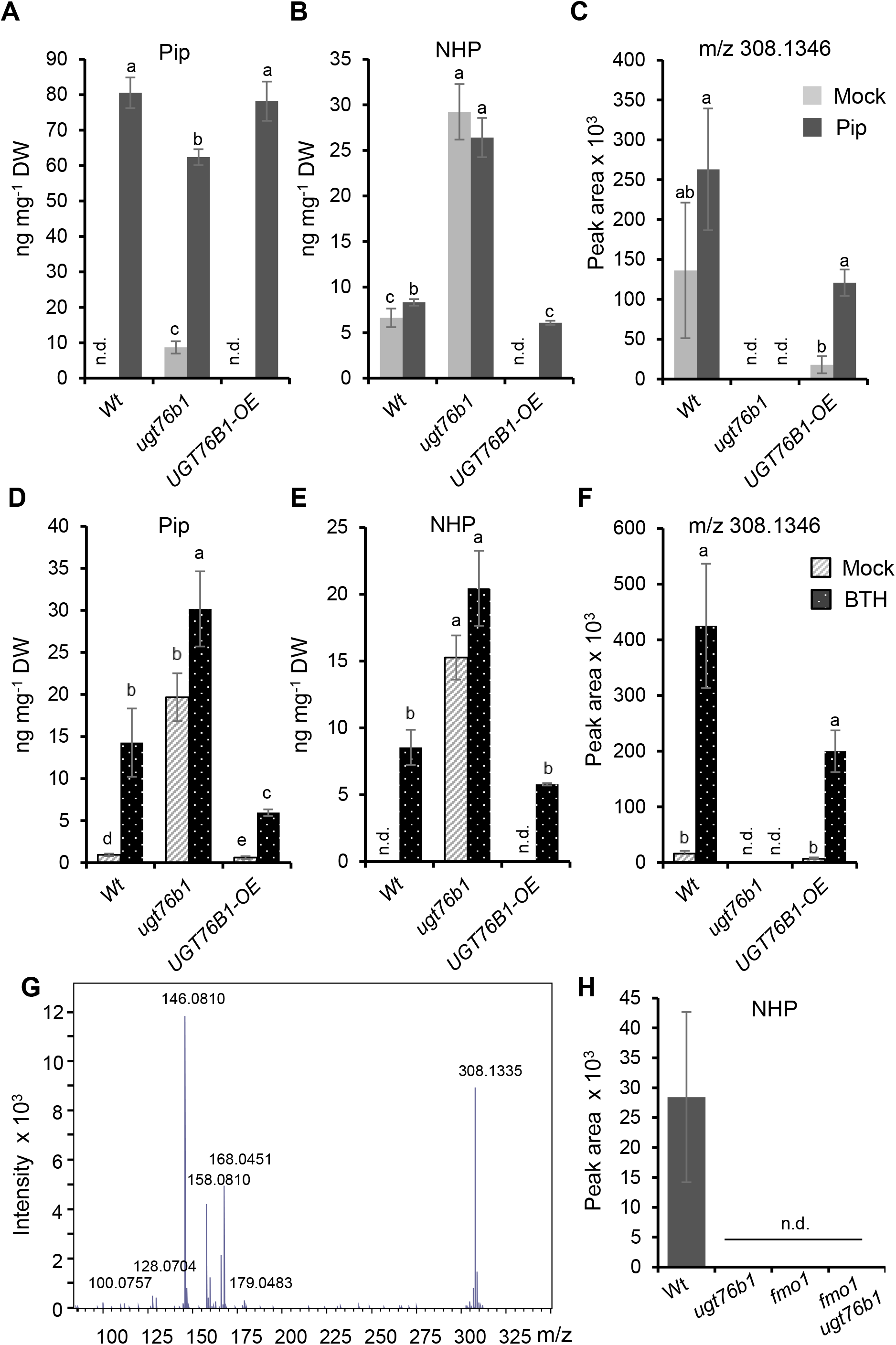
Accumulation of an NHP hexoside following treatment with resistance enhancers is dependent on UGT76B1. **(A)** to **(C)** Pip, NHP, and *m/z* 308.1346 content in rosette leaves of five-week-old wild type, *ugt76b1* and a UGT76B1-overexpressing line (OE) 48 h after watering with control solution or 1 mM DL-Pip. Rosette leaves were harvested, extracted and analyzed by LC-MS. **(D)** to **(F)** Detection of Pip, NHP, and *m/z* 308.1346 in leaf extracts of three-week-old plants sprayed with a control solution or 1 mM BTH. Leaves were harvested 48 h after the treatment and analyzed by LC-MS. Bars are means ± SD, n = 4; significant differences are indicated by letters according to one-way ANOVA (p.adj. < 0.05), n.d. = not detected. **(G)** MSMS fragmentation of *m/z* 308.1346 ± 0.005 identified in plant extracts. **(H)** Detection of *m/z* 308.1346 in extracts of wild-type Col-0 and in *ugt76b1*, *fmo1*, and *fmo1 ugt76b1* mutant lines using LC-MS (n.d., not detected).

To further investigate the role of UGT76B1 in defense-related metabolism, we comparatively analyzed extracts of wild-type, *ugt76b1*, and *UGT76B1*-OE plants treated with the priming-inducer benzothiadiazole (BTH) by LC-MS. The levels of Pip and NHP were induced more than tenfold in wild-type leaves 48 h after BTH treatment. The *UGT76B1*-OE line showed strongly reduced accumulation of Pip and NHP following BTH application, whereas the *ugt76b1* mutant over-accumulated both metabolites (Figure 1D and 1E). As observed above for the Pip treatments, the *ugt76b1* mutant plants were incapable of basal and BTH-induced formation of the putative NHP hexoside (Figure 1F). By contrast, the wild type and *UGT76B1*-OE plants substantially accumulated the NHP hexoside *m/z* 308.1346 after BTH application (Figure 1F).

To genetically substantiate that *m/z* 308.1346 is an NHP-derived hexoside, we further analyzed its occurrence in extracts of the *fmo1* mutant and an *fmo1 ugt76b1* double mutant. The *fmo1* background renders these lines incapable of NHP biosynthesis because of its defect in NHP-synthesizing FMO1 (Hartmann et al., 2018). Consistent with the proposed NHP hexoside structure, the *m/z* 308.1346 peak was undetectable in extracts of the *fmo1* and *fmo1 ugt76b1* lines (Fig. 1H). Thus, our comparative LC-MS metabolite analyses indicate that UGT76B1 mediates the Pip- and BTH-inducible formation of an NHP hexoside by Arabidopsis and thereby prevents over-accumulation of unconjugated NHP.

### Arabidopsis accumulates two NHP hexoside peaks in an FMO1-dependent manner

Previous GC-MS-based studies indicate that inoculation with a bacterial pathogen such as *Pseudomonas syringae* induces the formation of NHP and an NHP hexose conjugate (Hartmann et al., 2018; Hartmann and Zeier, 2018). In addition, an NHP hexoside with features similar to *m/z* 308.1346 (Figure 1G) was reported to accumulate in Arabidopsis in response to *P. syringae* attack (Chen et al., 2018). However, it was unresolved whether the NHP hexoside detected via LC-MS would correspond to the NHP hexose conjugate described by Hartmann and Zeier (2018) via GC-MS analyses, or whether two distinct NHP derivatives were identified in these studies.

To clarify this issue, we performed comparative metabolite analyses of leaf extracts from mock- and *P. syringae* pv. *maculicola* (*Psm*)-inoculated wild-type and *ugt76b1* plants using GC-MS analysis after trimethylsilylation of the extracted analytes according to previous work (Hartmann et al., 2018). Notably, we identified two putative NHP-hexoside peaks that both showed the *m/z* 172 mass spectral trimethylsilyl-N-hydroxypiperidine fragment characteristic for NHP. In addition, the mass spectra of both compounds displayed an *m/z* 652 in the higher spectrum range, which is consistent with a M^+^-CH_3_ ion of a penta-trimethylsilyated NHP-hexose conjugate, a thereof derived *m/z* 562 fragment, and further fragment ions characteristic for per-trimethlysilylated hexose conjugates (Figure 2A and 2B; Hartmann and Zeier, 2018). One of the two peaks present in the extracts from *Psm*-inoculated leaves was accumulating independently of UGT76B1. In fact, this peak represented the *Psm*-inducible NHP-hexoside previously described by Hartmann and Zeier (2018). Therefore, this substance was termed NHP-hexoside 1 (NHP-H1) (Figure 2A). In contrast, the second GC-MS-detected NHP-hexose conjugate, which we named NHP hexoside 2 (NHP-H2), was only detected in extracts from infected wild-type leaves, yet absent from *ugt76b1* mutant plants (Figure 2A and 2B). The UGT76B1-dependent accumulation of NHP-H2 by the GC-MS analyses was reminiscent of the accumulation pattern of the above-described *m/z* 308.1346 LC-MS peak, suggesting that NHP-H2 corresponds to the NHP hexoside identified by our LC-MS analyses (Figure 1). In contrast to the GC-MS analyses, a second UGT76B1-independent peak such as NHP-H1 was not unequivocally detected by our LC-MS settings.

**Figure 2.**
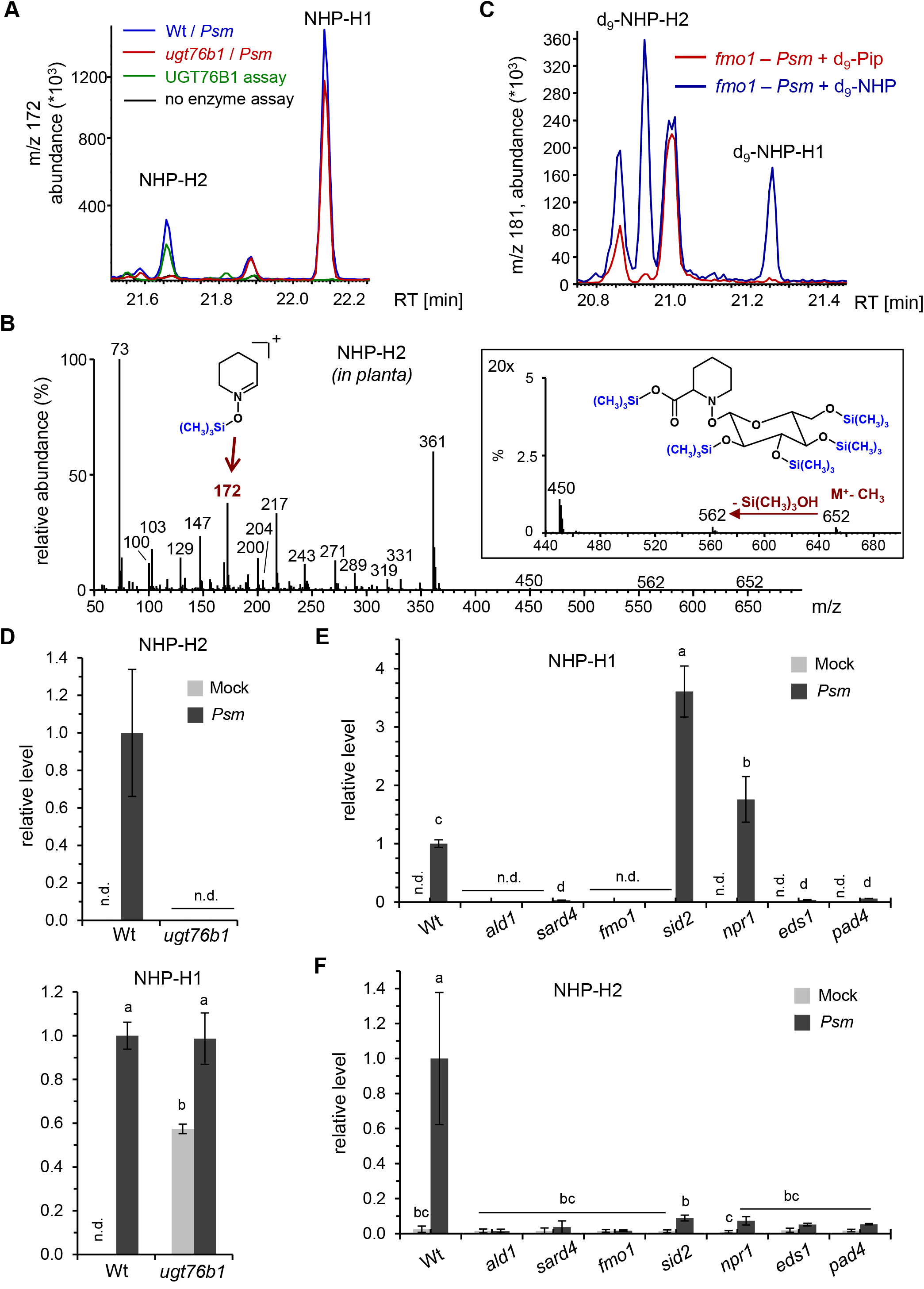
Inoculation with *Pseudomonas syringae* pv. *maculicola* induces accumulation of two distinct NHP hexoside derivatives dependent on a functional NHP biosynthetic pathway. **(A)** GC-MS analysis reveals the accumulation of two distinct NHP-hexose conjugates (NHP-H1 and NHP-H2) in leaves of *Psm*-inoculated wild-tpye plants. Overlaid ion chromatograms (*m/z* 172) are shown (blue: wild type, red: *ugt76b1*). NHP-H1 accumulates independently of UGT76B1 and represents the NHP-hexose-conjugate previously described by Hartmann and Zeier (2018). The *in planta* accumulation of NHP-H2 is absent in *ugt76b1* mutants. Moreover, NHP-H2 is synthesized *in vitro* by recombinant UGT76B1 (green: full enzyme assay with recombinant UGT76B1, black control assay without enzyme). Sample derivatisation by trimethylsilylation was performed prior to GC-MS analyses. **(B)** Mass spectrum of the penta-trimethylsilylated NHP-hexoside NHP-H2 (molecular weight: 667 g mol^−1^) from plant extracts. The M+-CH^3^ ion at *m/z* 652, which produces a *m/z* 562 ion by loss of Si(CH_3_)_3_OH, is clearly discernible. The structure of the trimethylsilyl-N-hydroxypiperidine fragment at *m/z* 172 is indicated. Fragment ions characteristic for per-trimethlysilylated hexose conjugates are *m/z* 450, *m/z* 361, *m/z* 271, *m/z* 217 and *m/z* 204 (Ehmann, 1974). Note that the mass spectrum of penta-trimethylsilylated NHP-H1 exhibits similar fragmentation patterns, including a more prominent *m/z* 172 (Hartmann and Zeier, 2018). **(C** Both NHP-H2 and NHP-H1 are biosynthetically derived from NHP. Feeding of deuterated D_9_-NHP to *Psm*-inoculated *fmo1* plants results, in contrast to feeding with D_9_-Pip, in the formation of D_9_-labelled NHP-H2 and D_9_-labelled NHP-H1. Ion chromatograms of *m/z* 181 are depicted. The *m/z* 181 ion corresponds to a D_9_-trimethlysilyl-hydroxypiperidine fragment. GC-MS analyses as described in (A) and (B). **(D)** Relative levels of NHP-H2 and NHP-H1 in *Psm*-inoculated or Mock-treated (MgCl_2_-infiltrated) leaves of wild-type and *ugt76b1* plants at 48 h post inoculation (hpi). Data represent the mean ± SD of three biological replicates (n.d., not detected). Mean levels in *Psm*-inoculated wild-type leaves are set to 1. Different letters denote significant differences (p < 0.05, ANOVA and post hoc Tukey HSD test). **(E)** and **(F)** Relative levels of NHP-H1 **(E)** and NHP-H2 **(F)** in *Psm*-inoculated or Mock-treated leaves of wild-type plants, NHP biosynthesis-defective mutants (*ald1, sard4, fmo1*), SA pathway mutants (*sid2, npr1*), as well as *eds1* and *pad4* mutants at 24 hpi. Other details as described in (D).

To substantiate the formation and identity of the two NHP hexosides by GC-MS, deuterium-labeled D_9_-Pip or D_9_-NHP was fed to *Psm*-inoculated *fmo1* plants. Both hexosides were absent from inoculated, D_9_-Pip-fed *fmo1* plants, indicating that their formation requires the FMO1-dependent Pip-to-NHP conversion (Figure 2C). Notably, deuterated NHP-H1 and NHP-H2 were detected in *fmo1* plants after D_9_-NHP feeding and subsequent *Psm* inoculation, corroborating their direct derivation from NHP *in planta* (Figure 2C). Moreover, the wild type accumulated both natural NHP hexosides in parallel with unbound NHP upon *Psm* infection, whereas they were virtually absent in non-inoculated control plants. However, NHP-H2 was UGT76B1-dependent and not detected in extracts of *Psm*-inoculated *ugt76b1* plants, whereas the level of NHP-H1 was similar in leaf extracts of *Psm*-inoculated wild-type and *ugt76b1* mutant plants (Figure 2D). Similar to the basal levels of free NHP, the basal levels of NHP-H1 were also strongly enhanced by naïve *ugt76b1* plants (Figure 2D). Together, our findings demonstrate the accumulation of two distinct NHP hexose derivatives upon pathogen inoculation, the UGT76B1-independent NHP-H1 and the UGT76B1-dependent NHP-H2.

We next examined the regulatory principles of the pathogen-inducible accumulation of these distinct NHP hexosides. Consistent with their biochemical derivation from NHP (Hartmann et al., 2018), the *Psm*-triggered accumulation of both NHP-H1 and NHP-H2 was absent in *ald1* and *fmo1*, and strongly diminished in *sard4* (Figure 2E and 2F). The accumulation of free NHP is also tightly regulated by the two immune-regulatory genes *ENHANCED DISEASE SUSCEPTIBILITY1 (EDS1)* and *PHYTOALEXIN-DEFICIENT4 (PAD4)* (Hartmann et al., 2018). A similar *EDS1*- and *PAD4*-dependency exists for the pathogen-induced biosynthesis of the two NHP hexosides, because the accumulation of both NHP-H1 and NHP-H2 was greatly suppressed by *eds1* and *pad4* mutant plants (Figure 2E and 2F). Previous analyses also revealed an over-accumulation of NHP by the Arabidopsis SA biosynthetic mutant *sid2* and by the SA-perception-defective mutant *npr1* during the course of *Psm* infection, indicating a modulation of NHP accumulation by an intact SA pathway (Hartmann et al., 2018). Interestingly, the accumulation of NHP-H2 was strongly dependent on intact *SID2* and *NPR1* genes, whereas NHP-H1 showed similar accumulation characteristics as free NHP and over-accumulated by *sid2* and *npr1* (Figure 2E and 2F). These analyses show that the inducible biosynthesis of the two identified NHP hexosides proceeds via an intact NHP biosynthetic pathway and, just like NHP biosynthesis, is boosted by the EDS1/PAD4-immune regulatory node. Notably, the pathogen-inducible generation of the two hexosides is oppositely regulated by SA signaling: whereas the accumulation of NHP-H1 and free NHP are negatively affected by a functional SA pathway, NHP-H2 accumulates in strong dependence of intact SA signaling.

### UGT76B1 glucosylates NHP to NHP-H2 *in vitro*

The complete loss of NHP-H2 by *ugt76b1* suggested that UGT76B1, known to glucosylate SA and isoleucic acid (ILA), may glycosylate NHP as an additional substrate. A homology model of UGT76B1 was created based on the crystal structure of the *Arabidopsis* SA glucosyltransferase UGT74F2 using the phyre2 software (Kelley et al., 2015; George Thompson et al., 2017). Interestingly, the predicted aglycon binding pocket of UGT76B1 is larger than that of UGT74F2 and, therefore, it is able to accommodate more spacious ligands. The *in silico* analysis predicts that NHP can alternatively bind to the active site of UGT76B1 in addition to the known aglyca SA and ILA (Figure 3A; Supplemental Figure 2).

**Figure 3.**
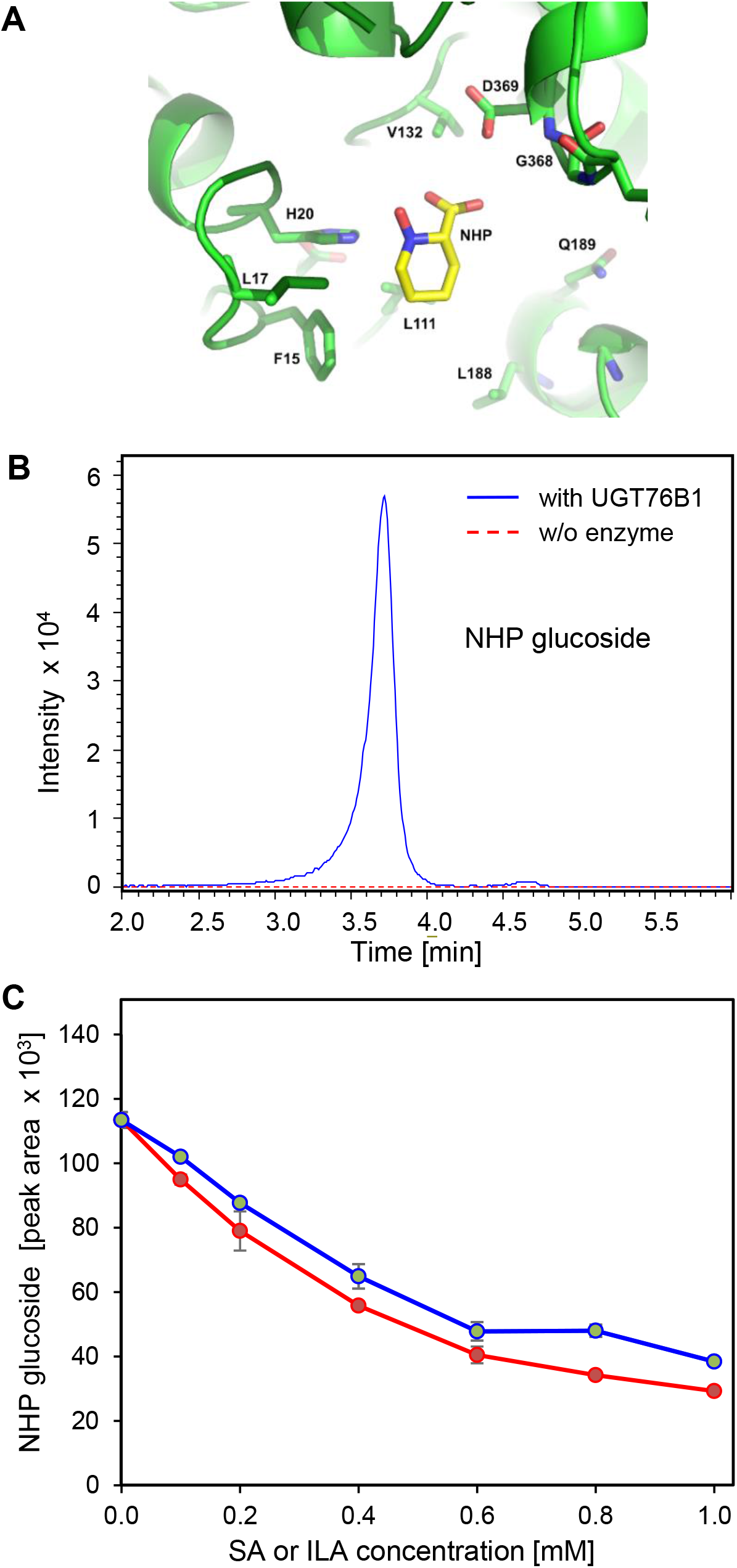
UGT76B1 glucosylates NHP *in vitro*, and its activity is inhibited by SA and ILA. **(A)** Computer modelling fitting NHP into the substrate binding pocket of UGT76B1 (Supplemental Figure 2). **(B)** LC-MS separation of the *in vitro* reaction of UGT76B1 (blue curve; control in stippled red) with NHP and UDP glucose producing an NHP glucoside with *m/z* 308.1346, which shows the correct MSMS fragmentation pattern (Supplemental Figure 1C). **(C)** Inhibition of UGT76B1-dependent NHP glucosylation by SA (red) and ILA (blue) *in vitro*. Values are means ± SE, n = 4.

The recombinantly produced UGT76B1 enzyme was incubated with NHP in the presence of UDP glucose to test whether it is able to catalyze the glucosylation of NHP *in vitro*. Indeed, LC-MS analysis of the enzymatic reactions revealed a peak with *m/z* 308.1346 co-eluting with the respective peak of plant extracts, whereas it was absent from the control incubation without the enzyme (Figure 3B; Supplemental Figure 1B). Fragmentation of *m/z* 308.1346 showed diagnostic fragments of an NHP hexoside, as observed for the metabolite from plant extracts (Figure 1G; Supplemental Figure 1C), supporting the formation of NHP glucoside by UGT76B1 *in vitro*. The NHP glucoside produced *in vitro* also co-eluted with the *in planta* detected NHP-H2 when analyzed by GC-MS, and showed the same mass spectral fragmentation pattern (Figure 2A and 2B; Supplemental Figure 3). Thus, the LC-MS feature *m/z* 308.1346 and the GC-MS peak NHP-H2 were coinciding in both analytical settings with the NHP glucoside produced *in vitro* by UGT76B1.

Maksym et al. (2018) had previously shown that ILA competitively inhibited the glucosylation of SA by UGT76B1. To investigate a possible influence of the two UGT76B1 co-substrates SA and ILA on UGT76B1-mediated NHP glucosylation, an *in vitro* competition assay was employed. The addition of increasing concentrations of ILA or SA from 0.1 mM to 1 mM steadily repressed the formation of NHP glucoside with a maximum inhibition of 65 and 75 % at 1 mM ILA and 1 mM SA, respectively (Figure 3C). Therefore, the UGT76B1-mediated glucosylation of NHP is competitively inhibited by the co-substrates SA and ILA.

Together, these analyses show that the UGT76B1 glycosyltransferase catalyzes the glucosylation of NHP *in vitro*. The *in vitro* reaction product is identical to the *in planta* detected, *UGT76B1*-dependent NHP-H2 hexoside. The usage of UDP-glucose as a substrate in our *in vitro* assays further reveals that NHP-H2 is an NHP glucose conjugate. The three UGT76B1 substrates SA, ILA, and NHP mutually compete with each other when analyzed with the glucosyltransferase *in vitro*, suggesting a regulatory interplay of these three immune-active metabolites via UGT76B1-dependent glucosylation.

### NHP-H1 and NHP-H2 are differentially conjugated hexosides

NHP possesses two reactive groups, a carboxylic acid and an N-OH hydroxyl group, which are amenable to sugar conjugation alternatively resulting in an ester bond and an *O*-β-glucoside, respectively. To get further insight into the nature of conjugation of NHP-H1 and NHP-H2, we employed hydrolytic digestions of leaf extracts with an esterase and a β-glucosidase. Besides NHP-H1 and NHP-H2, the extracts from *P. syringae*-inoculated wild-type leaves contained substantial levels of the SA glucose derivatives SA glucose ester (SGE) and SA-*O*-β-glucoside (SAG), which provided intrinsic ester- and *O*-β-glucoside-bound reference metabolites in this assay. All four hexose conjugates were stably detectable by GC-MS when incubated overnight with a sodium phosphate buffer of pH 6 (Figure 4). As expected, esterase strongly diminished the level of the reference metabolite SGE, whereas SAG remained unchanged. Consistent with previous digestion assays (Edwards, 1994), β-glucosidase treatment resulted in the partial cleavage of both the glucosidic bond of SAG and the ester linkage of SGE (Figure 4). Notably, NHP-H1 showed a similar sensitivity to the esterase- and β-glucosidase treatments as SGE, supporting a glucose ester structure for this conjugate (Figure 4). By contrast, NHP-H2, either extracted from wild-type leaves or synthesized *in vitro* by UGT76B1, showed resistance to esterase treatments (Figure 4 and Supplemental Figure 4), which was in line with the presumed identity of NHP-H2 as NHP-*O*-β-glucoside. However, β-glucosidase treatments did not diminish the NHP-H2 level either (Figure 4 and Supplemental Figure 4). This may be explained by a resistance of the special N-*O*-glucosidic linkage present in NHP-*O*-β-glucoside to the employed almond β-glucosidase, which chemically differs from the common C-*O*-glucosidic bond of SAG and other characterized β-glucosidase substrates. Together with previous mass spectrometric analyses of NHP-H1 (Hartmann and Zeier, 2018), these digestion assays support an NHP hexose ester structure for NHP-H1. In turn, the chromatographic and mass spectrometric distinction of the UGT76B1-synthesized NHP-H2 as well as its hydrolytic resistance suggest an N-O-glucosidic conjugation of NHP-H2 (Figure 2).

**Figure 4.**
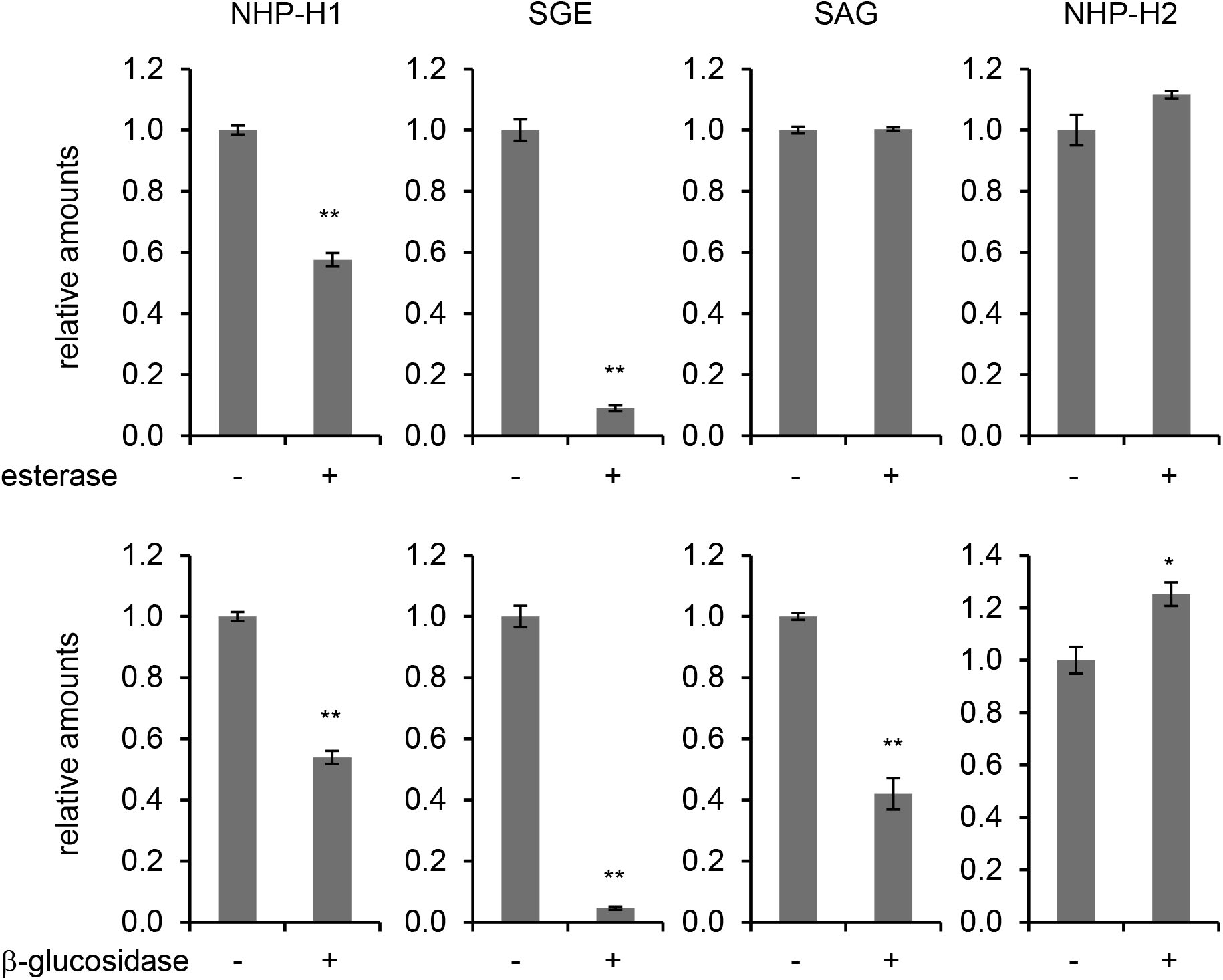
Effects of esterase and glucosidase treatments on NHP-H1, NHP-H2, salicylic acid glucose ester (SGE), and salicylic acid β-glucoside (SAG). An extract of Arabidopsis leaves inoculated with *P. syringae* pv. *maculicola* that accumulated NHP- and SA-conjugates was buffered to pH 6.0 with 0.1 mM sodium phosphate, and aliquots thereof incubated for 15 h with 10 U ml^−1^ esterase (+), 10 U ml^−1^ β-glucosidase (+), or with buffer only (-). Samples were analyses by GC-MS (Figure 2), and amounts of analytes were related to ribitol as internal standard. Values are expressed relative to the means of the buffer only condition (n = 3). Asterisks indicate significant differences between buffer only (-) and enzyme-treatments (** = p < 0.001 and * = p < 0.01 (two-tailed Student’s *t* test).

### Exogenous ILA promotes NHP accumulation

Exogenous ILA had previously been shown to induce defense responses and synergistically enhance the SA-dependent defense pathway (von Saint Paul et al., 2011; Bauer et al., 2020). After the identification of NHP as a third substrate of UGT76B1, we were interested to know how ILA would functionally interact with NHP and SA *in vivo*. Therefore, we treated wild-type plants with 500 μM ILA and determined the change in the levels of SA- and Pip-related metabolites. Interestingly, exogenously applied ILA significantly induced the amount of Pip and NHP 48 hours after the treatment (Figure 5A and 5B). In addition, NHP-H2 tended to be enhanced by the ILA treatment (Supplemental Figure 5A). To determine whether the NHP accumulation also involves the activation of the Pip biosynthetic pathway, we analyzed the expression of the NHP biosynthetic gene *FMO1*. Intriguingly, the expression of *FMO1* was induced by more than twofold at 48 hours after ILA application (Figure 5D). Similarly, we investigated the change in the abundance of SA-related metabolites after exogenous ILA application. SA, SAG, and SGE were enhanced by the treatment after 48 hours (Figure 5C; Supplemental Figure 5B and 5 C). Thus, ILA positively affects both the NHP and SA defense pathways suggesting a functional interaction between the UGT76B1 substrates ILA, SA and NHP in plant immunity.

**Figure 5.**
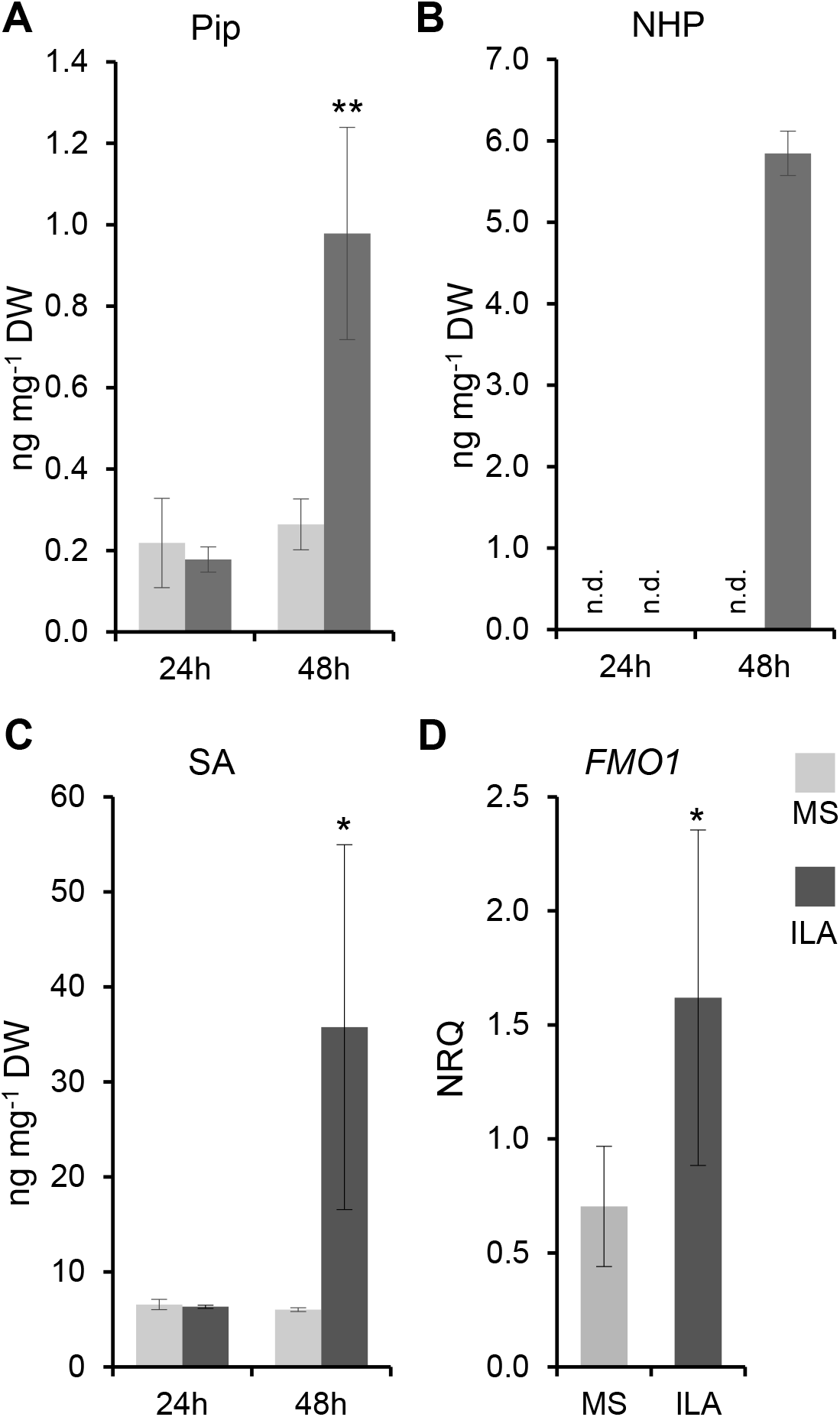
Exogenous ILA enhances the accumulation of SA, Pip, and NHP, and induces expression of *FMO1*. **(A)** to **(C)** Pip, NHP, and SA levels of leaves of 12-day-old wild-type seedlings 24 h and 48 h after incubation in ½ MS medium without (grey bar) and with 500 µM ILA (black bars). Bars represent means ± SD; n = 3-4. Differences between treated or untreated plants were analyzed by Welch two sample t-test; * = p < 0.05. **(D)** Transcript abundance of *FMO1* of leaves of 14-day-old plantlets grown in liquid culture were measured by RT-qPCR 48 h after the application of 500 µM ILA. Gene expression was normalized to S16 and UBQ5; bars are means ± SD; n = 4. Differences between treated or untreated plants were analysed by Welch two sample t-test; * = *p* < 0.05.

### Enhanced pathogen resistance of ugt76b1 requires intact FMO1

The *ugt76b1* mutant plants are more resistant to biotrophic pathogen attack than the wild type (von Saint Paul et al., 2011). This phenotype was related to the constitutively elevated level of SA and the upregulation of SA-dependent marker genes observed in this mutant, which, in turn, was attributed to the loss of UGT76B1 as an SA glucosyltransferase (von Saint Paul et al., 2011). However, the loss of *UGT76B1* also enhanced the levels of the defense-related metabolites Pip and NHP (Figure 1). Indeed, the NHP biosynthetic gene *FMO1* and the SA-biosynthetic gene *SID2* were constitutively upregulated in *ugt76b1* plants (Figure 6A; Supplemental Figure 6A). To examine the role of NHP in the enhanced resistance of *ugt76b1*, we employed the *fmo1 ugt76b1* double mutant line lacking NHP biosynthesis. Notably, the transcript abundances of *SID2* and of the SA-responsive genes *PR1* and *NPR1* were reduced to wild-type levels by the *fmo1 ugt76b1* double mutant (Supplemental Figure 6B and 6C). Furthermore, the enhanced levels of SA, SAG, SGE, and Pip found in *ugt76b1* leaves were reduced to wild-type levels by the introgression of *fmo1*, whereas the accumulation of NHP observed for *ugt76b1* was completely abolished by the *fmo1 ugt76b1* mutant background (Figure 6C to 6F). NHP-H2 was absent from all these mutants in accordance with the role of UGT76B1 and FMO1 in its biosynthesis (Figure 1H). Together, this indicates that the enhanced defense status of *ugt76b1*, which includes enhanced accumulation of the NHP precursor Pip and activation of the SA defense pathway, is triggered by the constitutive elevation of unconjugated NHP by *ugt76b1*.

**Figure 6.**
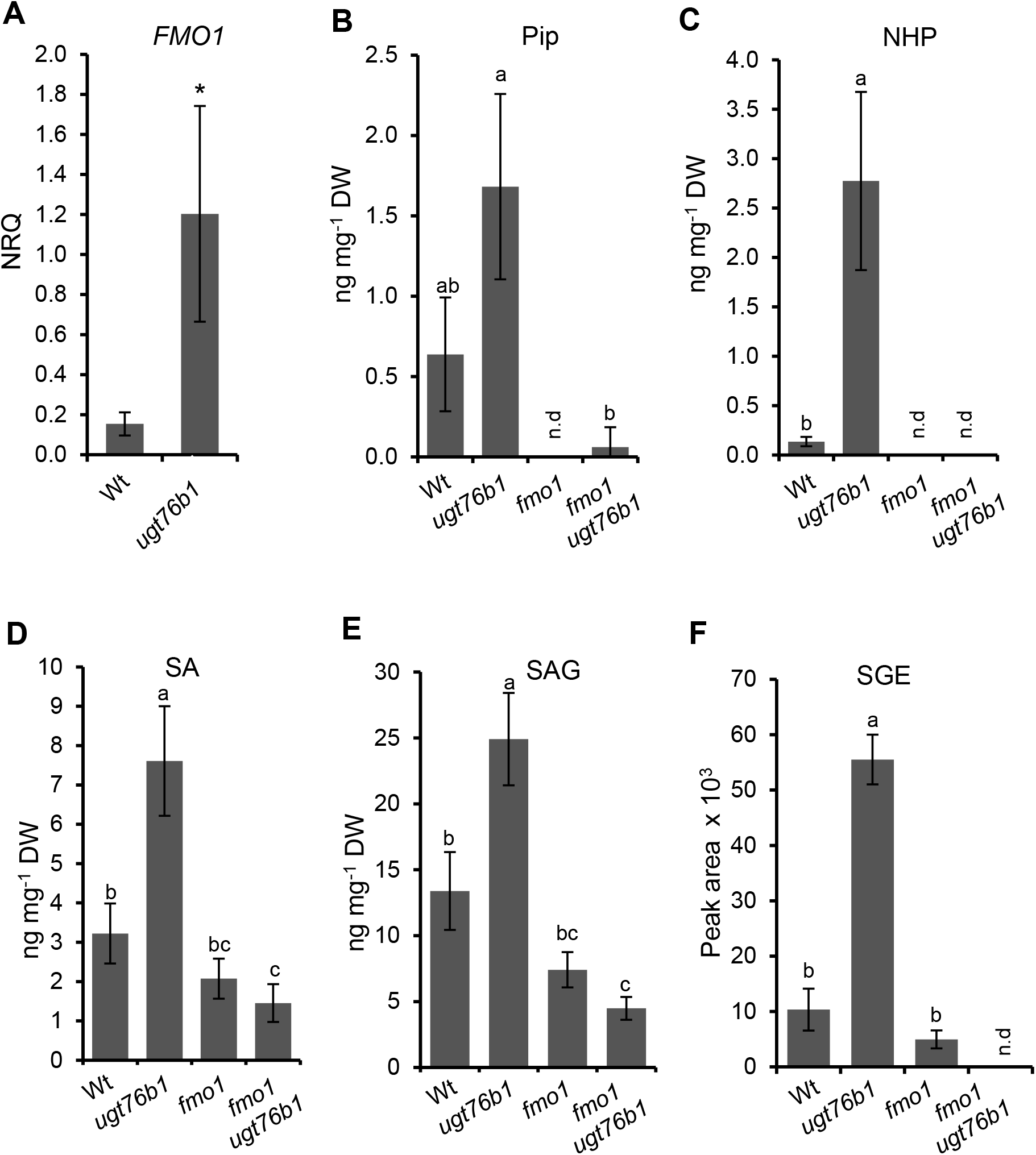
Enhanced accumulation of defense-related transcripts and metabolites by the *ugt76b1* mutant is dependent on FMO1. **(A)** RT-qPCR analysis of *FMO1* transcript abundance of leaves of four-week-old wild-type and *ugt76b1* plants grown under short day conditions; the normalized relative quantity (NRQ) was determined based on the *UBQ5* and *S16* internal standards; bars show means ± SD, n=4; differences between genotypes were analysed by Welch two sample t-test; * = *p* < 0.05. **(B)** to **(F)** Pip- and SA-related metabolites were determined using leaf extracts of four-week-old wild-type, *ugt76b1*, *fmo1* and *fmo1 ugt76b1* plants grown under short day conditions using LCMS. Bars show means ± SD, n = 4; differences between genotypes were analyzed by Welch two sample *t-test*; * = p < 0.05; n.d., not detected.

To directly examine the significance of accumulating NHP for the enhanced resistance phenotype of *ugt76b1*, we comparatively assayed basal resistance of wild-type, *fmo1*, *ugt76b1*, and *fmo1 ugt76b1* lines towards *Pseudomonas syringae* pv. *tomato* DC3000 (*Pst*). The loss of *FMO1* (and NHP) abolished the enhanced resistance phenotype of *ugt76b1*, since the *fmo1 ugt76b1* double mutant displayed a basal susceptibility similar to the *fmo1* single mutant or the wild type (Figure 7A). In parallel, the SA-depleted NahG *sid2* and NahG *sid2 ugt76b1* lines were employed to address the role of SA in the *ugt76b1*-associated resistance phenotype. The depletion of SA strongly raised the pathogen susceptibility of NahG *sid2* (Figure 7A). Notably, the introgression of the *ugt76b1* background into the NahG *sid2* line did not alter the resistance to bacterial infection, since NahG *sid2 ugt76b1* plants showed similar susceptibly to *Pst* as the SA-deficient NahG *sid2* line (Figure 7A). Together, these resistance assays indicate that the elevated NHP level of *ugt76b1* is the trigger for the primed defense status and enhanced basal resistance phenotype to bacterial infection. Elevated NHP activates SA biosynthesis in *ugt76b1*, and SA finally executes the NHP-initiated resistance response.

**Figure 7.**
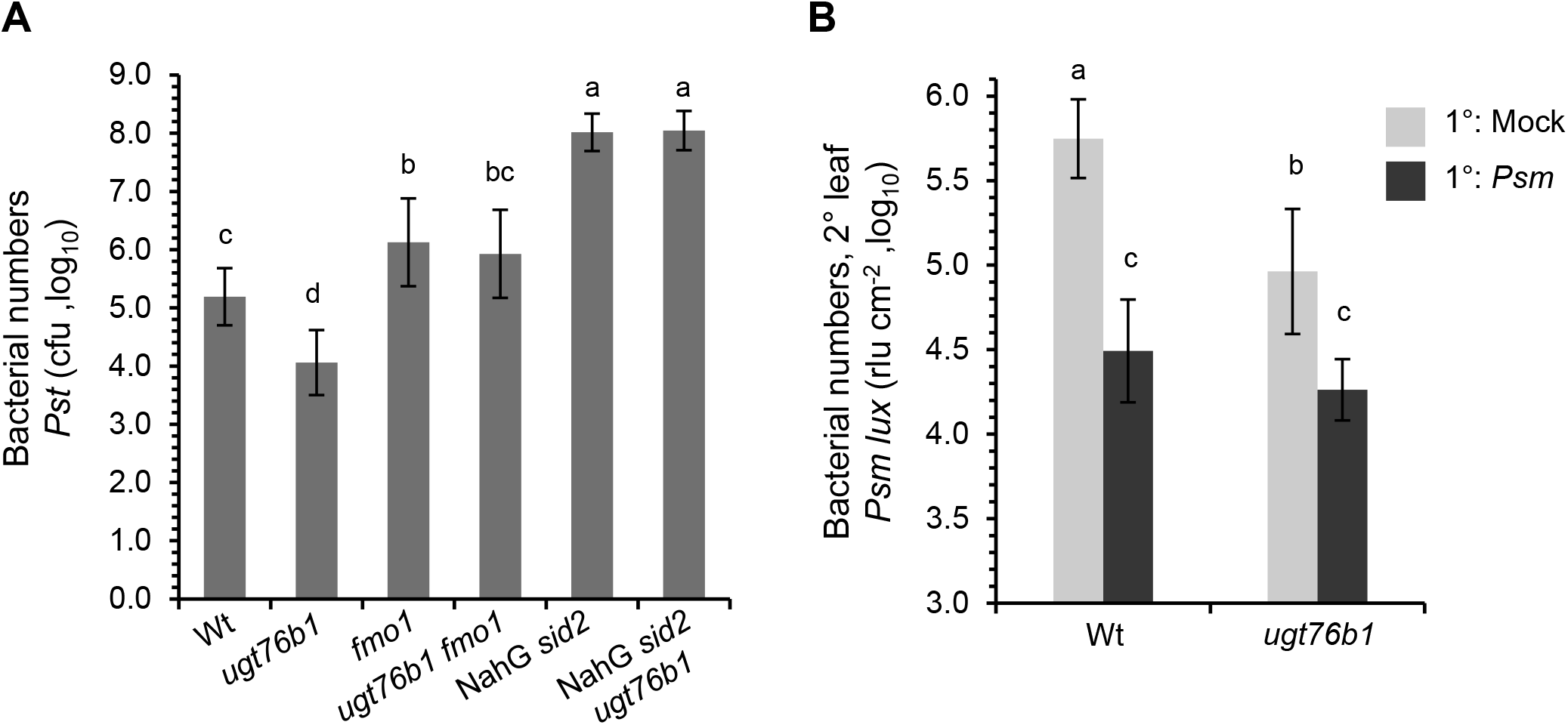
Local and systemic defense is enhanced by *ugt76b1* loss-of-function. **(A)** Susceptibility of *ugt76b1* introgressed into *fmo1* and NahG *sid2* towards *P. syringae* pv. *tomato* DC3000. Four-week-old wild-type (Wt), *ugt76b1*, *fmo1*, *fmo1 ugt76b1*, NahG *sid2* and NahG *sid2 ugt76b1* plants were infiltrated with 5 × 10^4^ cfu (OD_600_ = 0.0001) of *Pst* DC3000. Bacterial growth was monitored after 72 h. Bars are means ± SD of 15 replicates from three independent experiments, each experiment consisting of five biological replicates. The presented values are log10-transformed. Different letters denote significant differences (p < 0.01, ANOVA and post hoc Tukey HSD test). **(B)** The *ugt76b1* mutant is able to induce systemic acquired resistance. To assess SAR, three lower (1°) leaves of a plant were mock-treated or *Psm*-inoculated (OD_600_ = 0.005). Two days later, three upper (2°) leaves were challenge-infected with a bioluminescent *Psm* strain (*Psm* lux; OD_600_ = 0.001), and growth of *Psm* lux was assessed after 2.5 days by luminescence measurements. Bacterial numbers were determined as relative light units per leaf area (rlu cm^−2^). The presented values are log10-transformed. Bars are the mean ± SD of twelve or more replicate leaf samples from 6 to 7 different plants. Different letters denote significant differences (p < 0.01, ANOVA and post hoc Tukey HSD test). An independent experiment yielded similar results.

### SAR is still inducible by *ugt76b1*

NHP was previously identified as a main metabolic regulator of SAR of Arabidopsis (Chen et al., 2018; Hartmann et al., 2018). Therefore, we compared the SAR response of wild-type and *ugt76b1* mutant plants to examine whether the impact of UGT76B1 on NHP and the NHP glucoside NHP-H2 would affect SAR. Lower (1°) rosette leaves of a subset of plants were therefore pre-inoculated with *Psm* to induce SAR, whereas the 1° leaves of other plants were mock-treated for the examination of an appropriate non-induced control. Two days later, all the plants were challenge-infected with a bioluminescent *Psm* strain (*Psm* lux, Fan et al., 2008) and resistance to the bacterial infection was assessed by bioluminescence measurements 2.5 days later (Hartmann et al., 2017; Gruner et al., 2018). Inducing inoculations with *Psm* limited bacterial growth in challenged 2° leaves of the wild type compared to those of mock-control plants by about 16-fold, indicating a strong SAR establishment (Figure 7B). Moreover, mock-control plants of *ugt76b1* hosted significantly lower numbers of *Psm*-bacteria in their challenge-infected leaves than wild-type control plants, confirming that *ugt76b1* mutants exhibit enhanced basal resistance to bacterial infection (Figure 7B). Nevertheless, *ugt76b1* further enhanced resistance of 2° leaves upon an inducing 1° inoculation to resistance levels similar to those of the SAR-induced wild type (Figure 7B). Together, this indicates that naïve *ugt76b1* plants have per se acquired an enhanced state of pathogen resistance and, furthermore, are able to respond to pathogen inoculation by triggering a further SAR response. These resistance characteristics correlate with the enhanced basal levels and the *Psm*-induced accumulation of free NHP in *ugt76b1* (Figure 2).

## DISCUSSION

### UGT76B1 glucosylates NHP *in vivo* and *in vitro*

Previous *in vitro* and *in vivo* studies have shown that UGT76B1 catalyzes the glucosylation of SA and ILA (von Saint Paul et al., 2011; Noutoshi et al., 2012; Maksym et al., 2018; Bauer et al., 2020). Here, we identify NHP as a third substrate of the *Arabidopsis* UGT76B1 protein and provide evidence that UGT76B1 catalyzes the pathogen-inducible biosynthesis of NHP-*O*-β-glucoside, which we named NHP-H2 (Figure 2A; Figure 3). Arabidopsis leaves accumulate NHP-H2 in strict dependence of a functional *UGT76B1* gene upon inoculation with the bacterial pathogen *P. syringae*, in response to treatment with the priming-inducing chemical BTH and by exogenous feeding with Pip (Figure 1; Figure 2; Supplemental Figure 1). The accumulation of NHP-H2 by *Arabidopsis* further requires the NHP biosynthetic genes *ALD1*, *SARD4*, and *FMO1* in addition to *UGT76B1* (Figure 2F). The transcript levels of these four genes are strongly elevated upon microbial attack, indicating that the biosynthesis of NHP-H2 and its biosynthetic precursors NHP and Pip, is regulated at the level of transcription (Hartmann and Zeier, 2019). Previous and the present mutant analyses show that the immune regulators EDS1 and PAD4 positively influence Pip, NHP, and NHP hexoside accumulation (Figures 2E and 2F; Návarová et al., 2012; Hartmann et al., 2018). Moreover, the *in planta* conversion of D_9_-labelled NHP into D_9_-NHP-H2 corroborates that NHP is the direct biosynthetic precursor of the NHP glucoside, whereas D_9_-Pip was not able to support the formation of labeled NHP-H2 by the *fmo1* mutant (Figure 2C).

The current finding raises the number of substrates catalyzed by the UGT76B1 enzyme to three, *i.e.* SA, ILA, and NHP. This suggests a relatively low substrate specificity of the UGT76B1 glucosyltransferase. SA, ILA, and NHP share a carboxylic acid and a neighboring hydroxyl group as common features for the conjugation with the activated sugar molecule, although the planar structure of the aromatic SA differs from the aliphatic ILA and the heterocyclic NHP moieties. The structural modeling of the UGT76B1 protein shows the presence of a wide substrate-binding pocket due to the small amino acid side chains that face the catalytic site (Figure 3A; Supplemental Figure 2). Therefore, taking the size of the catalytic site and the structural similarity between SA and NHP into account, the modeling supports our experimental findings that SA, ILA, and NHP are alternative substrates of UGT76B1. Consequently, these substrates may compete with each other and thereby the reaction of one substrate may mutually inhibit the glucosylation of the respective other aglyca (Figure 8A). Indeed, the *in vitro* inhibition of SA glucosylation by ILA (Maksym et al., 2018) as well as the repression of NHP glucosylation by ILA or SA (Figure 3C) corroborates this metabolic interaction. Thereby, SA-, NHP-, and probably ILA-*O*-glucosides, but not the respective glucose esters are formed (Figure 4; see below; Li et al., 2015). Both ILA glucoside and NHP-H2 formation are strictly dependent on UGT76B1. In contrast, SAG can be also formed by other glucosyltransferases *in vitro* and *in vivo*, namely by AtUGT74F1 and AtUGT74F2 (Dean and Delaney, 2008; von Saint Paul et al., 2011; Noutoshi et al., 2012; George Thompson et al., 2017) (Figure 8A and 8B).

**Figure 8.**
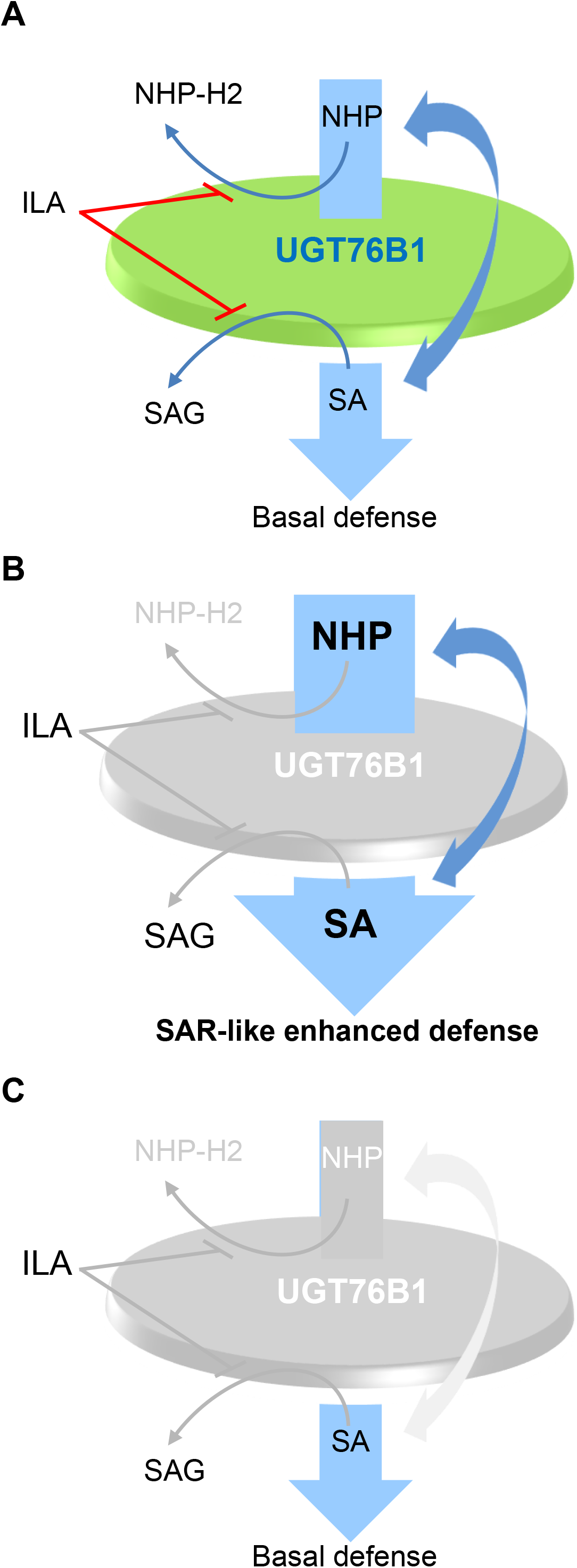
Model of UGT76B1 action towards NHP, ILA, and SA affecting basal defense. **(A)** UGT76B1 provides a metabolic hub controlling the levels of the unconjugated, immune-stimulating NHP and SA. Both substrates can be alternatively glucosylated by UGT76B1 to the putatively inactive SAG and NHP-H2. Thereby, the mutual amplification loops of NHP and SA and, consequently, basal defense are contained. ILA as an additional, competing substrate inhibits SAG and NHP-H2 formation and, therefore, enhances NHP- and SA-dependent defense. **(B)** The loss of UGT76B1 glucosylation releases this control. NHP and SA mutually enforced each other and promote an SA-dependent enhanced basal immune status. The loss of UGT76B1 abolishes the formation of ILA glucoside and NHP-H2, whereas SAG (and SGE, not included in model) is still produced due to the presence of other Arabidopsis SA glucosyltransferases, such as UGT74F1. In addition, the elevated NHP level is accompanied by the enhanced formation to the UGT76B1-independent NHP-H1 (not included in the model). **(C)** The additional loss of FMO1 reverses the primed immune status of *ugt76b1* to a basal defense level. Thus, the enhanced defense scenario observed in (B) is dependent on NHP and similar to SAR. In the absence of FMO1 and UGT76B1 the regulatory metabolic hub is completely eliminated.

### The UGT76B1-dependent glucoside NHP-H2 is different from the putative NHP glucose ester NHP-H1

NHP contains an N-OH and a carboxylic hydroxyl group. Basically, both groups can be conjugated to glucose, which would result in the formation of an NHP-*O*-glucoside and an NHP glucose ester, respectively. This is reminiscent to the conjugation modes of SA, for which the *O*-glucoside SAG and the glucose ester SGE exist (Dean and Delaney, 2008; George Thompson et al., 2017). The comparison of LC-MS and GC-MS analyses revealed that the UGT76B1-dependent *O*-glucoside NHP-H2 was not identical with a previously identified NHP hexoside (Hartmann and Zeier, 2018), consequently termed NHP-H1. The GC-MS-derived mass spectra of the GC-separable silylated NHP-H1 and NHP-H2 peaks and labelling assays using D_9_-NHP support their annotation as NHP-containing hexose conjugates (Figures 2B and 2C; Ehmann, 1974; Hartmann et al., 2018). Notably, the NHP-characterizing *m/z* 172 trimethylsilyl-N-hydroxypiperidine ion occurs more prominently in the NHP-H1 than in the NHP-H2 spectrum. This supports an NHP glucose ester structure for NHP-H1 and an NHP-*O*-glucoside structure for NHP-H2, since the glucose ester can be directly cleaved into *m/z* 172, whereas the glucoside can only produce this fragment by an additional rearrangement. The NHP glucose ester structure of NHP-H1 is further supported by our esterase and β-glucosidase digestion assays. NHP-H1 was digested in a similar manner as the SA glucose ester SGE, whereas NHP-H2 was completely esterase-insensitive (Figure 4; Supplemental Figure 4). However, unlike SAG, NHP-H2 also showed resistance to β-glucosidase treatments (Figure 4 and Supplemental Figure 4), which might be due to the unusual N-*O*-glucosidic bond present in an NHP-*O*-β-glucoside. Nevertheless, these analyses and the known biochemistry of UGT76B1 catalyzing *O*-β-glucoside formation of ILA and SA clearly support an NHP glucoside structure of NHP-H2 (Figure 1; Supplemental Figure 1). Based on the LC-MS-derived characteristics (Figure 1; Supplemental Figure 1), NHP-H2 may be identical to the previously LC-MS-detected NHP hexoside described by Chen and colleagues (2018).

Interestingly, the two identified NHP hexose derivatives differ with respect to the regulation of their biosynthesis by SA: the accumulation of NHP-H1 is negatively influenced by an intact SA pathway, whereas NHP-H2 generation requires intact SA biosynthesis (ICS1/SID2) and perception (NPR1). Furthermore, NHP-H1 also hyper-accumulates in conjunction with the *ugt76b1-*enhanced basal defense and shows an accumulation pattern similar to free NHP (Figures 2E and 2F; Hartmann et al., 2018). The reduced accumulation of NHP-H2 by SA pathway mutants could also be the result of a reduced expression of *UGT76B1*. Indeed, the transcript abundance of *UGT76B1* was repressed by the *sid2* mutant following mock- and *Psm-*treatments (Bernsdorff et al., 2016). Thus, the over-accumulation of NHP and NHP-H1 by SA pathway-defective mutants may result from an attenuating NHP-H2 production. However, the functional implications of these different regulatory patterns are not clear yet. Moreover, the glycosyltransferase responsible for the biosynthesis of NHP-H1 and the biological significance of NHP-H1 formation remain to be determined.

### UGT76B1 modulates free NHP levels and plant acquired resistance

Unstressed plants keep NHP at very low basal levels and exhibit only modest basal immunity against infection by adapted pathogens (Figure 1; Figure 7; Hartmann et al., 2018). Upon pathogen contact, however, wild-type plants can induce SAR to subsequent infection and thereby achieve a primed state of enhanced systemic immunity that allows a boosted activation of defense responses (Fu and Dong, 2013; Shah and Zeier, 2013). A series of studies demonstrates that the pathogen-induced activation of the pipecolate pathway, which entails accumulation of Pip and NHP in locally inoculated and distant leaf tissue, is a key and indispensable event to trigger SAR and the associated primed state (Návarová et al., 2012; Bernsdorff et al., 2016; Chen et al., 2018; Hartmann et al., 2018). The decisive trigger that switches on SAR of wild-type plants is the endogenous accumulation of NHP in response to an inducing pathogen attack. Exogenous feeding of plants with NHP enhances immunity in a similar manner as pathogen-induced SAR, indicating that elevations in the levels of NHP are sufficient for the induction of acquired resistance (Chen et al., 2018; Hartmann et al., 2018).

Our results show that the *ugt76b1* mutant markedly elevates the endogenous level of NHP already in the absence of pathogen contact. This goes hand in hand with enhanced basal resistance to *P. syringae* and increased basal levels of SA, SA derivatives, and defense-related gene expression of this mutant (Figure 1; Figure 6; Figure 7; Supplemental Figure 6). Therefore, the *ugt76b1* mutant exhibits a constitutively primed, SAR-like state (Figure 8A and 8B). However, *ugt76b1* plants also show a reduced rosette size under normal growth conditions as a negative impact (von St. Paul et al., 2011). Notably, basal resistance, SA accumulation, and defense-related gene expression of the *ugt76b1 fmo1* double mutant, which is entirely blocked in NHP biosynthesis because of its defect in the Pip hydroxylase FMO1, is similar to or even below wild-type like levels (Figure 1; Figure 6; Figure 7; Supplemental Figure 6; Figure 8C). This indicates that elevated NHP triggers immune priming of *ugt76b1*. Thus, UGT76B1 as an SA, ILA, and, in addition, NHP glucosyltransferase suppresses such an NHP-activated and SA-dependent defense status of wild-type plants (Figure 8; see below).

Two recently elucidated examples further illustrate that genetically de-regulated immune signaling pathways of Arabidopsis can result in the constitutive activation of plant NHP biosynthesis, and consequently, augmented immunity as well as growth impairment. Mutational defects in the transcription factor CAMTA3, a negative regulator of NHP and SA biosynthetic genes, resulted in elevated basal NHP levels, autoimmunity and dwarfism. Both the constitutive activation of immunity and the dwarfed phenotype were suppressed by the NHP-deficient *ald1* and *fmo1* backgrounds (Kim et al., 2020; Sun et al., 2020). Moreover, overexpression of the Arabidopsis calcium-dependent protein kinase CPK5 resulted in basal NHP accumulation, enhanced pathogen resistance and growth defects, all of which were dependent on functional *FMO1* (Guerra et al., 2020). Thus, the de-regulated overproduction of NHP as well as its impaired biochemical inactivation, as observed by *ugt76b1*, can cause resistance-enhancing phenotypes and a disturbed balance of growth and defense.

### UGT76B1 functions as a regulatory hub integrating the immune-active small molecules NHP, ILA, and SA

UGT76B1 glucosylates ILA and SA in addition to its newly identified activity towards NHP. SA is known as an important plant stress hormone that is crucial for basal resistance, effector-triggered immunity, and SAR (Klessig et al., 2018). Therefore, UGT76B1-dependent SA glucosylation is an important means to negatively control these responses, in conjunction with at least two other SA glucosyltransferases of Arabidopsis, AtUGT74F1 and AtUGT74F2 (Dean and Delaney, 2008; von Saint Paul et al., 2011; Noutoshi et al., 2012; George Thompson et al., 2017). In addition, plants unable to accumulate SA because of expression of the bacterial SA hydroxylase NahG or defects in endogenous SA biosynthesis display strongly impaired local resistance to diverse virulent and avirulent pathogens and are compromised in SAR (Gaffney et al., 1993; Volko et al., 1998; Nawrath and Métraux, 1999; Bernsdorff et al., 2016). Moreover, NHP will only be able to induce a strong SAR, if plants possess an intact SA biosynthesis pathway, indicating a close cooperation and mutual activation of NHP and SA in SAR induction (Bernsdorff et al., 2016; Hartmann et al., 2018; Figure 8). Consistently, the *ugt76b1* mutation that elevates free NHP levels does no longer cause induction of acquired resistance when introgressed to the SA-depleted NahG *sid2* background. In fact, NahG *sid2* and NahG *sid2 ugt76b1* plants display a similar and strongly attenuated resistance to *P. syringae*, which is more severely compromised than the resistance of *fmo1* and *ugt76b1 fmo1* lines (Figure 7A; Figure 8C). This illustrates the key function of SA for both basal immunity and acquired resistance and indicates that SA is the final executor of the NHP-triggered resistance response observed by *ugt76b1* (Figure 8). However, the SA pathway can be also activated independently of NHP at pathogen inoculation sites, as shown for *Pseudomonas*-inoculated *ald1* mutants devoid of Pip and NHP (Bernsdorff et al., 2016). This scenario implies that there are additional interactions regulating NHP and SA levels.

ILA is a further player of the UGT76B1-associated immune hub. Exogenous application of ILA activates defense in a synergistic manner with SA (Bauer et al., 2020) and it increases NHP and SA levels (Figure 5). *In vitro*, ILA inhibits the UGT76B1-mediated glucosylation of NHP and SA, which may contribute to the increased levels of both immune modulators *in vivo* (Figure 3; Figure 8A; Maksym et al., 2018). The *ugt76b1* mutant is also defective in ILA glucosylation and exhibits increased endogenous ILA levels (von Saint Paul et al., 2011; Maksym et al., 2018), which, in turn, may also contribute to the primed state of *ugt76b1*. However, ILA may not enhance defense apart from its UGT76B1-mediated action, because its exogenous application to *ugt76b1* mutants did not amplify SA-induced expression of defense marker genes (Bauer et al., 2020; Figure 8B). On the other hand, ILA may play a role in mitigating defense responses. At the sites of *Pst* inoculation, the endogenous ILA concentration is lowered in response to infection (Maksym et al., 2018; Bauer et al., 2020). Thus, the ILA-mediated inhibition of NHP and SA glucosylation by UGT76B1 would be diminished. Consequently, an enhanced glucosylation of NHP and SA would attenuate the defense signaling towards a balanced immune status in the course of infection. Currently, these roles of ILA cannot be ultimately resolved *in vivo*, since the biosynthetic pathway and corresponding loss-of-function mutants have not yet been identified.

In conclusion, UGT76B1 provides a common regulatory hub orchestrating basal defense by the glucosylation of the immune-active small molecules NHP, ILA, and SA. NHP and SA foster defense and mutually amplify each other, while their action is further promoted by ILA (Figure 8; Bernsdorff et al., 2016; Chen et al., 2018; Wang et al., 2018; Bauer et al., 2020). The conjugation by UGT76B1 contains the levels of active NHP and SA in an interactive manner due to the mutual competition of the aglycon substrates. The loss of functional UGT76B1 releases this control and leads to an activated state of basal defense, which is triggered by NHP, further enhanced by ILA, and eventually executed by SA as the central downstream signal. Thus, *ugt76b1* establishes a SAR-like status in the absence of a pathogen infestation (Figure 8). In contrast, the promiscuous glucosylating activity of functional UGT76B1 maintains a non-activated, steady state of basal defense of naïve plants and may enable the dynamic, interactive control of the immune-active compounds after a pathogen infection.

## METHODS

### Plant materials and growth conditions

Wild-type *Arabidopsis thaliana* accession Columbia (WT), *ugt76b1-1* knockout mutant (SAIL_1171A11), a line constitutively overexpressing UGT76B1 (von Saint Paul et al., 2011), *fmo1-1* (SALK_026163; Bartsch et al., 2006; Mishina and Zeier, 2006), the SA-depleted double mutant NahG *sid2* (Veeragoni et al., 2020), and NahG *sid2 ugt76b1* triple mutants were used. *fmo1 ugt76b1* was generated from the above single mutants and provided by Wei Zhang. Multiple mutants were obtained by genetic crossing. Mutants were obtained from the *Arabidopsis* seed stock center (Scholl et al., 2000) except for NahG and *sid2-1* (Gaffney et al., 1993; Nawrath and Métraux, 1999). Plants were grown in a controlled growth chamber (light / dark regime 10 / 14 h at 20 / 16 °C, 80 / 65 % relative humidity, light at 130 µmol m^−2^ s^−1^) on a peat moss-based substrate (Floragard Multiplication substrate, Oldenburg, Germany) : quartz sand (8 : 1) mixture. For liquid cultures, ½ MS medium with 1% sucrose was used and the seedling were germinated and grown while shaking at 100 rpm in the same conditions described above.

### Application of chemicals and sample collection

Benzothiadiazole (BTH; Syngenta) was applied by spraying a 1 mM aqueous solution onto three-week-old plants; more than twenty rosettes were harvested after 24 or 48 h. For pipecolic acid treatment five-week-old plants were either watered with 10 ml water or 1 mM D/L-pipecolic acid (Pip; Sigma-Aldrich) solution as described before (Návarová et al., 2012); leaves of five rosettes were harvested 48 h after the onset of the treatment for one biological sample. For ILA (Interchim) treatment, approximately twenty wild type seedlings were germinated and grown for 12 days in six-well plates under conditions described above. Then, the media of one half of the samples was replaced with fresh ½ MS and the remaining half with 500 μM ILA containing ½ MS medium. The treatment was applied for 24 h and 48 h. Finally, the shoots were harvested after washing with distilled water and blotting on tissue paper. In all cases, the harvested samples were immediately frozen in liquid nitrogen.

### In planta labeling experiments with isotope-labeled metabolites

Treatments with isotope-labeled metabolites were essentially performed as described in Hartmann et al., 2018. Three to four mature leaves of 5-week-old soil-grown Arabidopsis wildtype or *fmo1* plants were infiltrated in the morning either with *Psm* (OD_600_ = 0.005) or 10 MgCl_2_ (Mock) as described above. Four to six hours after the inoculation event, the same leaves were infiltrated with a 2 mM solution of d_9_-labelled NHP prepared freshly in HPLC-grade water. Water infiltrations served as control treatments. DL-2-piperidine-d_9_ carboxylic acid (D_9_-Pip; Sigma-Aldrich 688444) was co-infiltrated at a final concentration of 2 mM as part of the final bacterial suspension or Mock-solution. Infiltrated leaves were harvested at 48 hpi (in reference to the first infiltration event with *Psm* or mock solutions) and GC-MS analysis of trimethylsilyl-derivatized metabolites was carried out according to the described analytical protocols (section: GC-MS analyses). Authentic d_9_-labeled 1-hydroxypiperidine-2-carboxylic acid (d_9_-N-hydroxypipecolic acid, d_9_-NHP) was synthesized according to a protocol of Murahashi and Shiota (1987), as described in Hartmann et al. (2018) using piperidine-d_11_ (CAS Number: 143317-90-2) as reactant.

### UGT enzyme assay

Purification of recombinant UGT76B1 and enzyme activity testing were performed as described (Meßner et al., 2003) using 0.5 µg recombinant GST-UGT fusion protein, 0.5 mM NHP and 2 mM UDP glucose in a 50 µl reaction volume at 30°C for 30 min. The product formation was analysed by LC-MS. For enzyme competition assays, the reaction mixture contained 0.5 mM SA and varying concentrations of NHP or ILA. The formation of SAG was measured by absorption at 302 nm after HPLC separation (Meßner et al., 2003). Preparative reactions to produce NHP-H2 using UGT76B1 and SGE-containing SA glucosides using UGT74F2 were incubated at least 2 h.

### Esterase and β-glucosidase digestion

Leaf extracts from 5-week-old *P. syringae* pv. *maculicola* ES4326 (*Psm*)-inoculated Arabidopsis Col-0 plants (48 hpi) accumulating the NHP- and SA-conjugates of interest were used for esterase and β-glucosidase digestion experiments. Shock-frozen leaf tissue was freshly-ground to a fine powder and extracted twice with ice-cold MeOH/H_2_O (80:20, v/v) in a ratio of 1 ml per 100 mg leaf tissue. The resulting extract was then evaporated to dryness and re-suspended in water. Aliquots of this solution were incubated in sodium phosphate buffer (50 mM final, pH 6.0) for 15 h at 30°C, either in the presence of 50 u ml^−1^ porcine liver esterase (E3019, Sigma-Aldrich) or 0.5 u ml^−1^ β-glucosidase from almonds (7512.1, Carl Roth). Aliquots incubated in buffer without the respective enzymes served as control treatments. Reactions were stopped by addition of an excess amount of MeOH/H_2_O solution (80:20, v/v) that was supplemented with an internal standard mix. Samples were then immediately analysed according to the GC-MS-based analytical procedure described above. The amounts of NHP-H1, NHP-H2, SAG, and SGE were quantified in relation to ribitol (CAS N°: 488-81-3, A5502 Sigma-Aldrich) as internal standard.

The hydrolytic experiments were independently repeated using the *in vitro* produced NHP-H2 and an NHP-H2 containing leaf extracts (prepared from four-week-old wild-type plants spray-treated with 1 mM BTH for 48 h) using LC-MS analytics. Aliquots of the *in vitro* reaction or freeze-dried extracts dissolved in water were treated with 15 u ml^−1^ porcine liver esterase (pH 8.0) or 0.9 u ml^−1^ almond β-glucosidase (pH 5.0) (E2884, G0395; Sigma-Aldrich) for 5 h incubation at 30 °C. SAG and SGE contained in the extracts were used as internal controls of the hydrolysis by glucosidase and esterase; in fact, SGE was even susceptible to the alkaline condition (pH 8) used for esterase incubation.

### LC-MS analyses

For determination of SA- and Pip-related metabolites, frozen-materials were crushed to powder and lyophilized. The extraction and measurement of the compounds was carried out as described before with minor modifications (Wenig et al., 2019). Briefly, ~20 mg of freeze-dried material was resuspended in 1.5 ml 70% methanol, vigorously shaken at 4 °C for 1 h and subsequently centrifuged at 18.000 × g for 10 min. Supernatants were then transferred to new 2-ml tubes, concentrated *in vacuo* (RVC 2-25 CDplus, Christ) for 2 h and shortly frozen at −80 °C. The samples were freeze-dried overnight, re-dissolved in 100 µl acetonitrile: H_2_O (1:1, v/v) and then centrifuged for 5 min at 18.000 *g* at 4 °C. Next, 90 µl of supernatants were transferred onto microwell plates fitted with 0.2 µM PVDF filters and centrifuged. Five µl of the extract was injected twice as technical replicate for the LC-MS analyses. Pip, NHP, and NHP-H2 were detected in positive ionization mode, whereas SA, SAG, and SGE were measured in negative ionization mode. Mass spectra were acquired in a mass range of 50-1300 m/z. SA (Sigma-Aldrich), SAG (Santa Cruz Biotechnology), SGE, Pip (Sigma-Aldrich), and NHP were identified using authentic standards. SGE was produced using recombinant *Arabidopsis* UGT74F2 expressed as a GST fusion protein (Lim et al., 2002; Meßner et al., 2003). NHP was synthesized according to Hartmann et al. (2018) and confirmed by GC-MS analyses or by LC mass spectrometry. LC-MS/MS fragmentation patterns of NHP and NHP-H2 were in accordance with Chen et al. (2018).

SA and SAG were quantified against an internal standard curve with ten calibration points and two internal standards (p-nitrophenol, Fluka; camphorsulfonic acid, Sigma-Aldrich) at 1 mg l^-1^. Pip and NHP were quantified against external standard curves of (D,L-piperidine-2-carboxylic acid, Sigma-Aldrich) and NHP with six calibration points. Normalized peak areas were used as a semi-quantitative estimate of NHP-H2 and SGE, since the reference compounds were obtained as unpurified reaction products of the recombinant glucosyltransferases UGT76B1 and UGT74F2, respectively. SGE peaks were normalized with the internal standard camphorsulfonic acid. Normalization of Pip, NHP, and NHP-H2 was performed using the total ion chromatogram during the gradient elution. Retention times and *m/z* values: SAG 8.0-8.5 min, 299.0750; SGE 9.2-9.5 min, 299.0750; SA 10.0-10.2 min, 137.0250; internal standards p-nitrophenol 10.0-10.1 min, 138.0195; camphorsulfonic acid 9.0 min, 231.0695; Pip 2.1-2.4 min, 130.0860; NHP 1.4-1.7 min, 146.0817; NHP-H2: 3.3-3.7 min, 308.1346.

### GC-MS analyses

Gas chromatography-mass spectrometry (GC-MS)-based analyses of plant metabolites (NHP-H1, NHP-H2, SAG, SGE) was performed as detailed in Hartmann et al. (2018) and Stahl et al. (2019) with minor modifications. Briefly, 50 mg of pulverized, frozen leaf samples were extracted twice with 1 ml of MeOH/H2O (80:20, v/v). For internal standardization, 1 µg of salicin and ribitol were added. 600 μl of the extract were evaporated to dryness, re-dissolved in 100 µl of 100 mM sodium phosphate (pH 6.0), and the solvent removed again. Free hydroxyl groups of the analytes were converted into their trimethylsilyl derivatives by adding 20 µl of pyridine, 20 µl of N-methyl-N-trimethylsilyltrifluoroacetamide (MSTFA) containing 1% trimethylchlorosilane (v/v) and 60 µl of hexane. The mixture was heated to 70 °C for 30 min. After cooling, samples were diluted with hexane, and 2 µl of the solution was separated on a gas chromatograph (GC 7890A; Agilent Technologies) equipped with a Phenomenex ZB-35 (30m × 0.25mm × 0.25µm) capillary column. The following GC temperature settings were used: 70 °C for 2 min, with 10 °C / min to 320 °C, 320 °C for 5 min. Mass spectra were recorded in the electron ionization (EI) mode between m/z 50 and m/z 750 with a 5975C mass spectrometric detector (Agilent Technologies). Metabolites were analyzed using the Agilent MSD ChemStation software. For quantification, substance peaks of selected ion chromatograms (NHP-H1: *m/z* 172; NHP-H2: *m/z* 172; SAG: *m/z* 267; SGE: *m/z* 193) were integrated, and peak areas of plant metabolites were related to those of the internal standards salicin *(m/z* 268) or ribitol (*m/z* 307).

### RT-qPCR

For determination of the expression of defense related genes, leaf samples were collected from soil grown four-week-old wild type, *ugt76b1*, *fmo1* and *fmo1 ugt76b1* lines grown under conditions described above. Rosettes of three plants were pooled to represent one biological sample and each genotype was analyzed in four replicates. For determination of *FMO1* expression of ILA-treated wild-type seedlings, treatments and sample collection was carried out as described above. RNA extraction, cDNA synthesis and expression analysis of the genes was performed according to Bauer et al. (2020).

### Modeling

A homology model of UGT76B1 was created on the template of UGT74F2 with SA and UDP glucose (PDB ID: 5U6M, George Thompson et al., 2017) using the *phyre2* software (http://www.sbg.bio.ic.ac.uk/phyre2; Kelley et al., 2015) with manual adjustments. The position of UDP-glucose in the structure was deduced by superposition of UGT76B1 model with flavonoid 3-O glucosyltransferase (PDB ID: 2C1Z; Offen et al., 2006). The 3D models of N-OH Pip, SA and ILA acid were placed into the protein using the docking program *PatchDock* (https://bioinfo3d.cs.tau.ac.il/PatchDock/index.html) and analyzed using *COOT* (Emsley et al., 2010). The localization near UDP-glucose was used as an additional criterion for positioning the aglycon substrates.

### Bacterial inoculation and infection assays

Full-grown leaves of four-to five-week-old wild-type, *ugt76b1*, *fmo1*, *fmo1 ugt76b1*, NahG *sid2* and NahG *sid2 ugt76b1* plants were gently pressure infiltrated either with 10 mM MgCl_2_ as control, or with suspensions of *Pseudomonas syringae* pv. *tomato* DC3000 (*Pst*) or *Pseudomonas syringae* pv. *maculicola* strain ES4326 (*Psm;* Hartmann et al., 2017) in 10 mM MgCl_2_. For basal resistance assays, plants inoculated with *Pst* (OD_600nm_ = 0.0001) were grown for three days (72 h) at growth conditions described above. Three leaf discs from three independent plants, representing one biological sample, were pooled 2 h and 72 h after the inoculation and immersed in 500 µl of 10 mM MgCl_2_ solution containing 0.01% Silwet L70 (Lehle Seeds). The analysis of the bacterial growth was carried out as described before (Katagari et al., 2002). Each treatment was replicated five times and the complete experiment was independently repeated two times. Combined data of all experiments are displayed. For the determination of metabolite accumulation in response to *Psm*, a bacterial suspension of OD_600nm_ = 0.005 was used for leaf inoculation. Mock- and *Psm*-inoculated leaves were harvested at different times after the treatment. Six leaves from two different plants were routinely pooled for one biological replicate, and three to four biological replicates per treatment analyzed by GC-MS.

### SAR assays

To induce SAR, three lower (1°) rosette leaves of five-week-old, short-day-grown Arabidopsis plants were inoculated with *Psm* by gently syringe-infiltrating bacterial suspensions in 10 mM MgCl_2_ (OD_600_ = 0.005) into the abaxial sides of the leaves (Gruner et al., 2018). Analogous infiltration of a 10 mM MgCl_2_ solution served as a mock-control treatment. Two days later, three upper (2°) rosette leaves of the plants were challenge-inoculated by infiltration with a suspension (OD_600_ = 0.001) of *Psm* carrying the *Photorhabdus luminescens luxCDABE* operon (*Psm* lux; Fan et al., 2008). Bacterial growth was assessed 2.5 d later by punching leaf discs from inoculated leaf areas and determining the bioluminescence of *Psm* lux with a Sirius FB12 luminometer (Berthold Detection Systems). Bacterial numbers were expressed as relative light units (rlu) per cm^2^ leaf area, which show strong positively correlations with colony-forming units (cfu cm^−2^) assessed by the more traditional, plating-based assays (Gruner et al., 2018).

### Statistics

Statistical analyses were performed in R (R 3.5.1 for Windows; https://www.r-project.org/). To test for statistical differences in datasets of bacterial growth experiments (Figure 7) and GC-MS-based metabolite measurements (Figure 2), log_10_-transformed measuring values were subject to ANOVA and a post-hoc Tukey’s HSD test as detailed previously (Hartmann et al., 2018). For robust statistical analyses of other data, the WRS2 package based on Wilcox’ WRS functions was used. Two groups were compared via Welch two sample t-tests. One-way multiple group comparisons were tested in R using the robust one-way ANOVA function *t1way* with lincon post hoc test. p-values were Holm-corrected and adjusted p-values were used for analysis (Mair & Wilcox, 2019).

## Supplemental Data

**Supplemental Figure 1.** UGT76B1 loss-of-function abolished the occurrence of m/z 308.1246 that co-elutes with the *in vitro* produced NHP glucoside.

**Supplemental Figure 3.** The GC/MS-mass spectra of NHP-H2 detected *in vitro* and *in planta* are identical.

**Supplemental Figure 2.** Structural modelling of aglyca at the substrate binding pocket of UGT76B1.

**Supplemental Figure 4.** *In vitro* synthesized NHP glucoside as well as the *in vivo* NHP-H2 peaks show resistance to esterase and β-glucosidase treatments.

**Supplemental Figure 5.** Exogenous ILA enhances the accumulation of NHP-H2, SAG, and SGE.

**Supplemental Figure 6**. Introgression of *fmo1* mutation into *ugt76b1* reverses the expression of *SID2* and defense marker genes to wild-type level.

## ACKNOWLEDGEMENTS

We are grateful to Wei Zhang for discussion, critical reading of the manuscript and generation of the *fmo1 ugt76b1* double mutant. Elisabeth Georgii supported us by discussion and statistical analyses. Corina Vlot-Schuster and Jörg Durner were instrumental throughout the project. The work was supported by the Helmholtz Zentrum München (A.R.S) and a grant to J.Z. by the German Research Foundation (DFG grant ZE467/6-2). Finally, we greatly acknowledge the exchange of data with Jianghua Cai and Asaph Aharoni about their independent finding that UGT76B1 glucosylates NHP.

## AUTHOR CONTRIBUTIONS

D.W.M., S.B., and M.H. performed experiments. M.H. and J.Z. analyzed metabolites by GC-MS, B.G. contributed expression analyses, and B.L. performed LC-MS experiments. R.J. modelled the binding of substrates at the active site. D.W.M., S.B., M.H., J.Z., and A.R.S. conceived the project and designed experiments; D.W.M., S.B., J.Z., and A.R.S. wrote the paper. All authors read and approved the final version of the article.

Submission: June 19, 2020

## Notes

### Competing Interest Statement

The authors have declared no competing interest.

