## Supplemental Figures for "UGT76B1, a promiscuous hub of small molecule-based immune signaling, glucosylates N-hydroxypipecolic acid and controls basal pathogen defense"

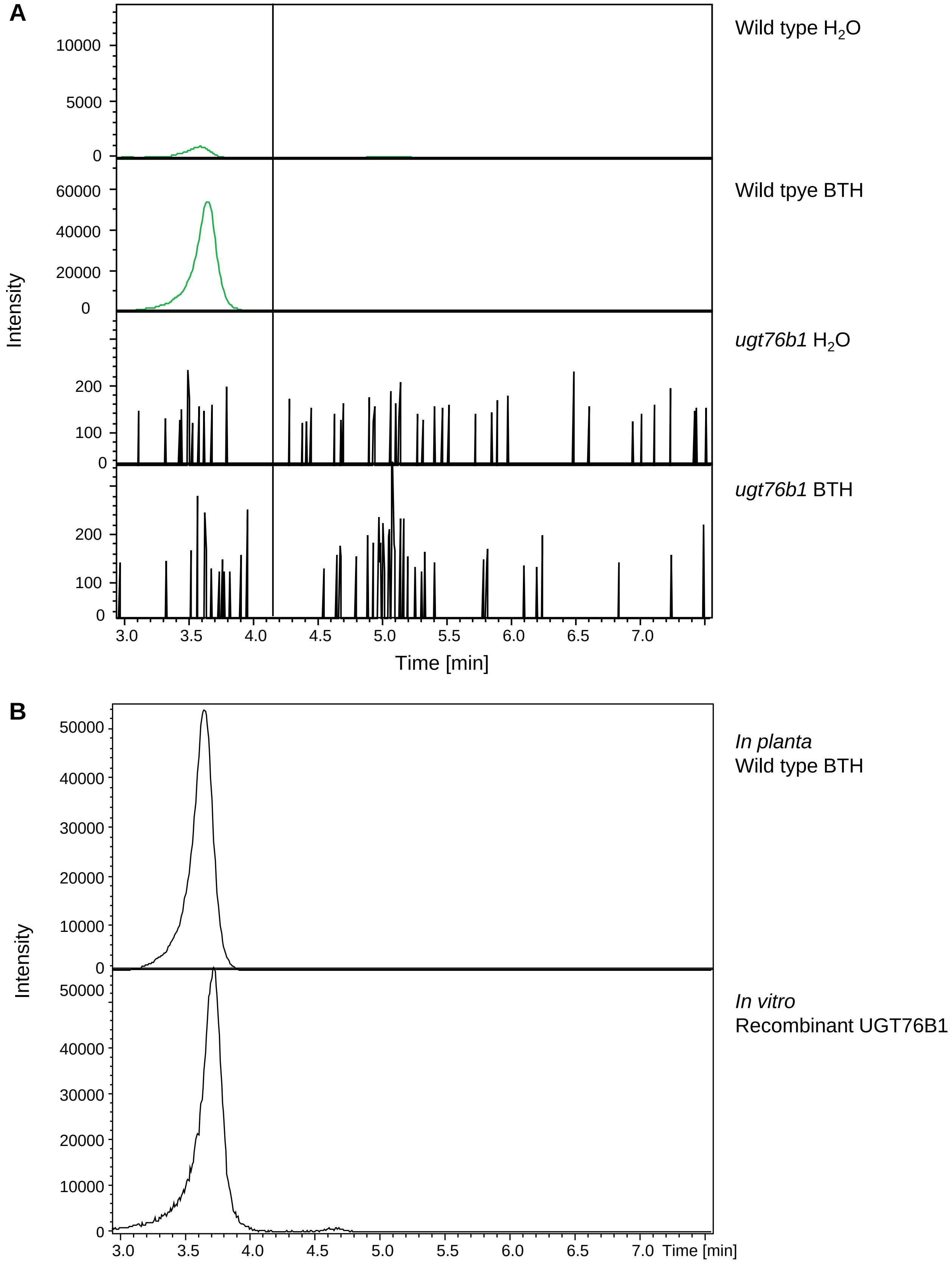


**Supplemental Figure 1 (continued on next page)**

**C**

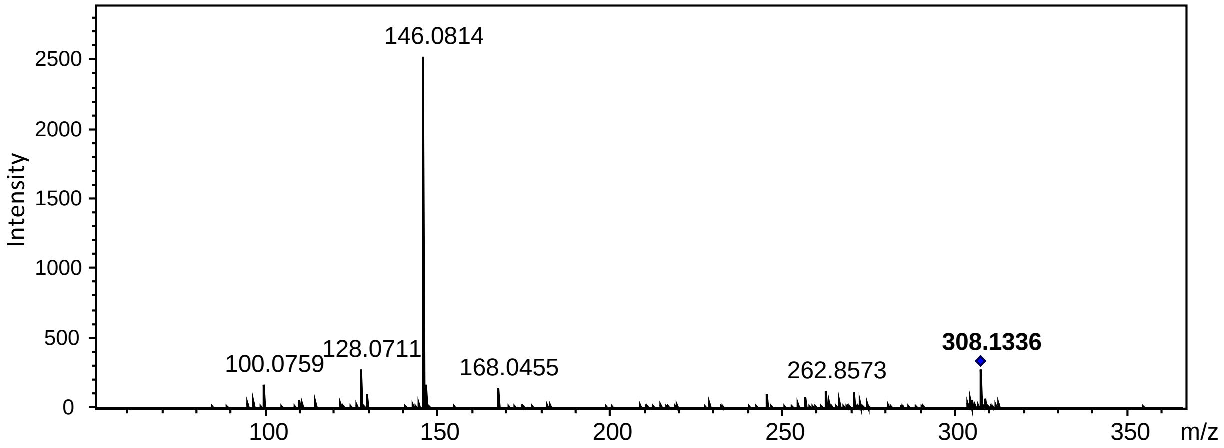


**Supplemental Figure 1.** UGT76B1 loss-of-function abolished the occurrence of *m/z* 308.1246 that co-elutes with the *in vitro* produced NHP glucoside.

**(A)** The *in vivo* metabolite peak *m/z* 308.1346 is enhanced by BTH treatment and dependent on UGT76B1.

**(B)** Co-elution of *m/z* 308.1346 peaks of plant extract and *in vitro* product of recombinant UGT76B1.

**(C)** LC-MSMS analysis and fragmentation of the *m/z* 308.1346 peak of the *in vitro* reaction of recombinant UGT76B1 using NHP and UDP glucose as substrates.

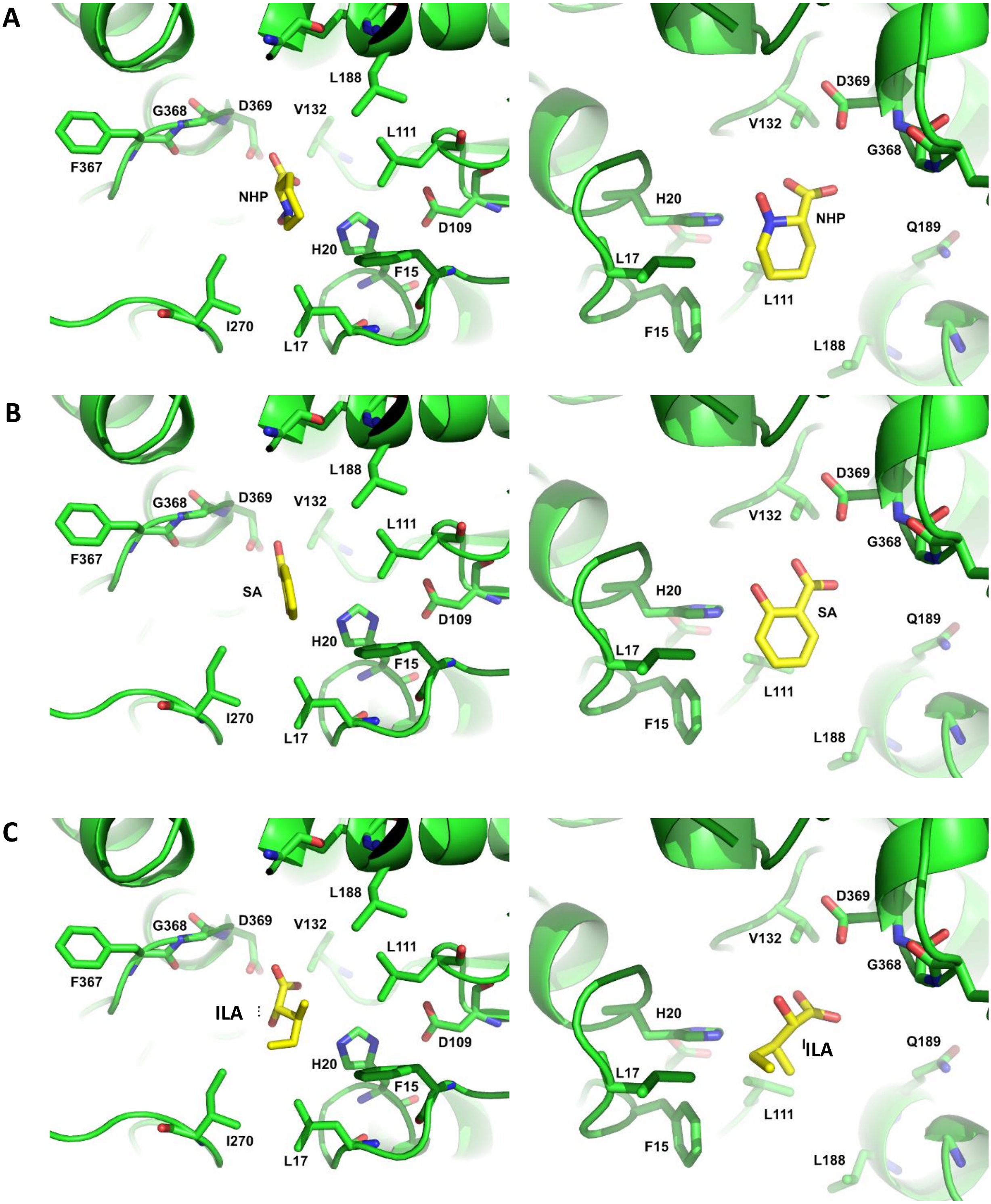


**Supplemental Figure 2.** Structural modeling of aglyca at the substrate binding pocket of UGT76B1.

**(A)** NHP, **(B)** SA, and **(C)** ILA binding is shown from two separate views each. The orientation towards the catalytic side chains (George Thompson et al., 2017) is chosen to enable the formation of an *O*-glucosidic bond.

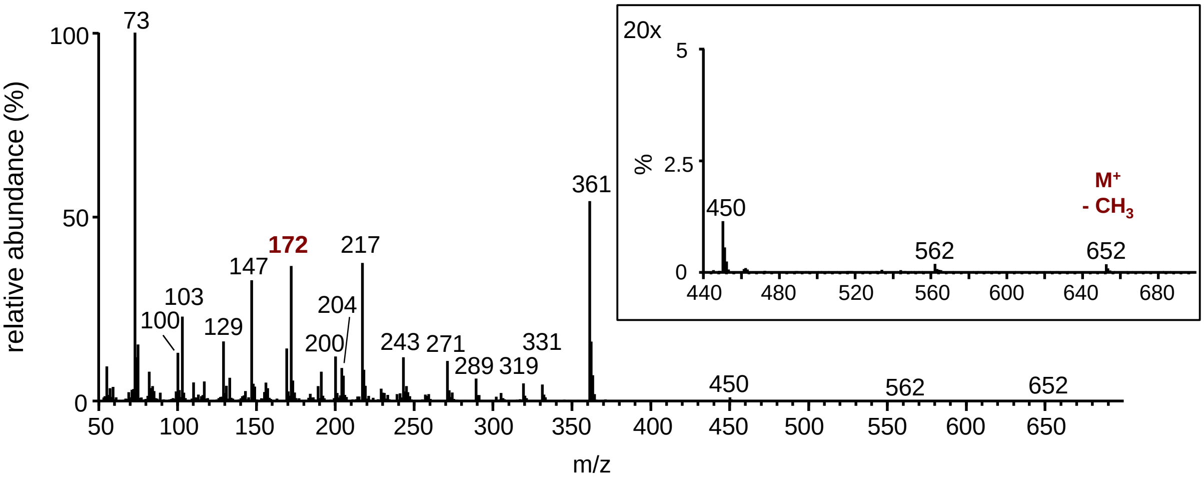


**Supplemental Figure 3.** The GC/MS-mass spectra of NHP-H2 detected *in vitro* and *in planta* are identical.

Mass spectrum of the per-trimethylsilylated NHP glucoside NHP-H2 (molecular weight = 667 g $mol^{-1}$) produced by the recombinant UGT76B1, as obtained by GC-MS analysis. Note that the spectrum is virtually identical to the spectrum of the in *planta*-detected NHP-H2 (Fig. 2C). Moreover, the GC-MS retention times of *in vitro* and *in planta* detected NHP-H2 are identical (Fig. 2B). The $M^+$-$CH_3$ ion (*m/z* 652) is clearly discernible. The *m/z* 172 ion, which occurs as the main fragment in the mass spectrum of per-trimethylsilylated NHP-H1 (Hartmann and Zeier, 2018), is consistent with a trimethlysilylated hydroxypiperidine fragment.

A

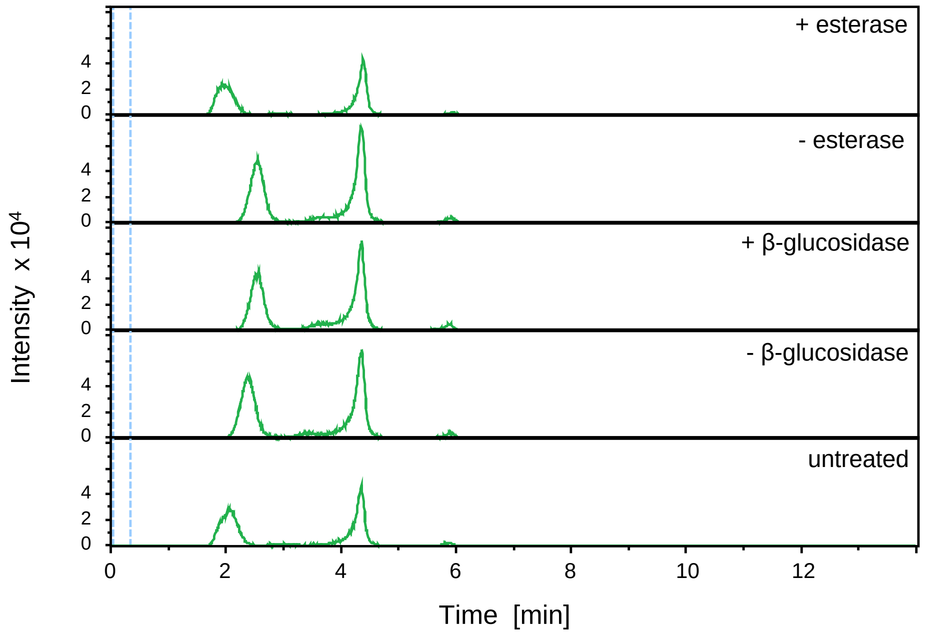


B

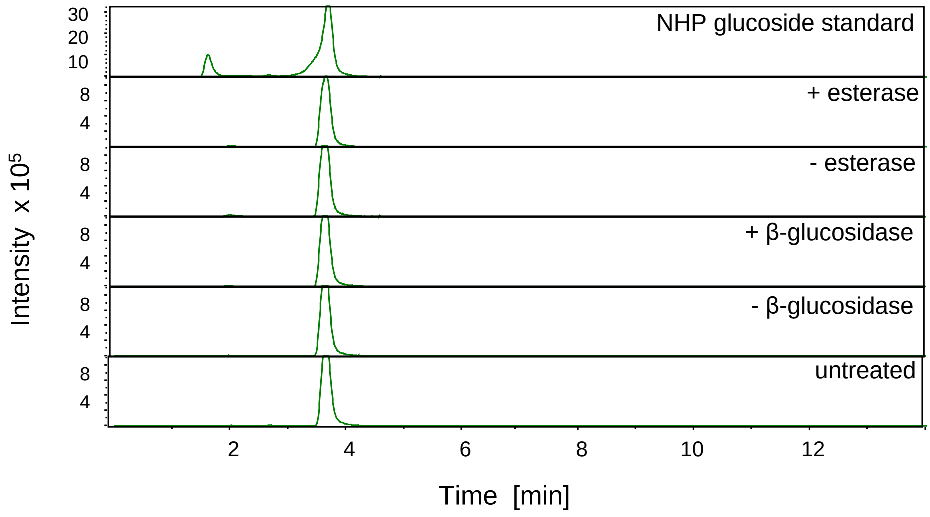


**Supplemental Figure 4 (continued on next page)**

C

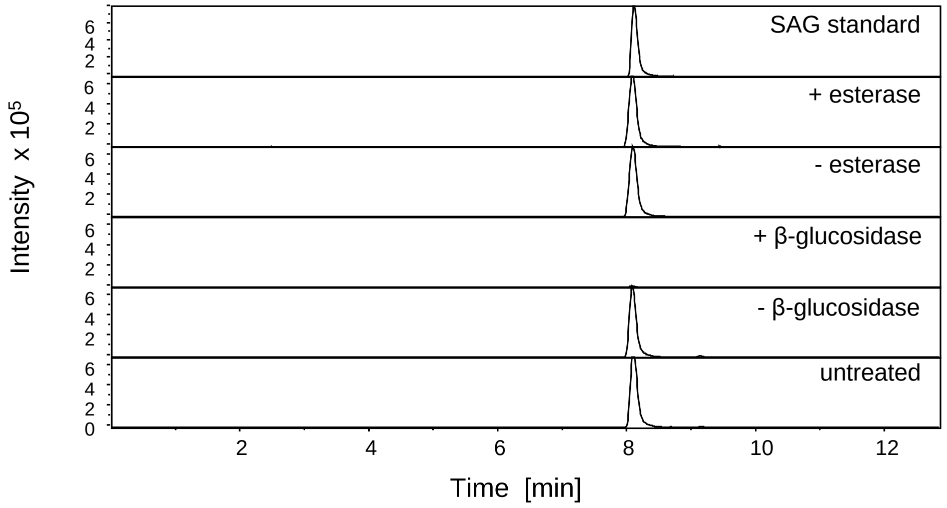


D

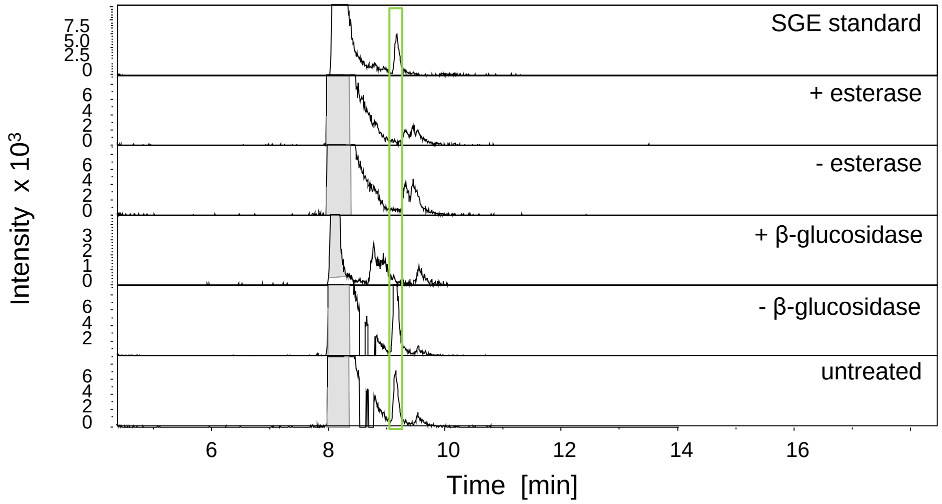


**Supplemental Figure 4.** *In vitro* synthesized NHP glucoside as well as the *in vivo* NHP-H2 peaks show resistance to esterase and β-glucosidase treatments.

**(A)** *In vitro* product NHP glucoside +/- esterase, +/- ß-glucosidase.

**(B)** Plant m/z 308.1346 +/- esterase, +/- ß-glucosidase.

**(C)** Plant extract-contained SAG +/- esterase, +/- ß-glucosidase.

**(D)** Plant extract-contained SGE (highlighted small peak) +/- esterase, +/- ß-glucosidase

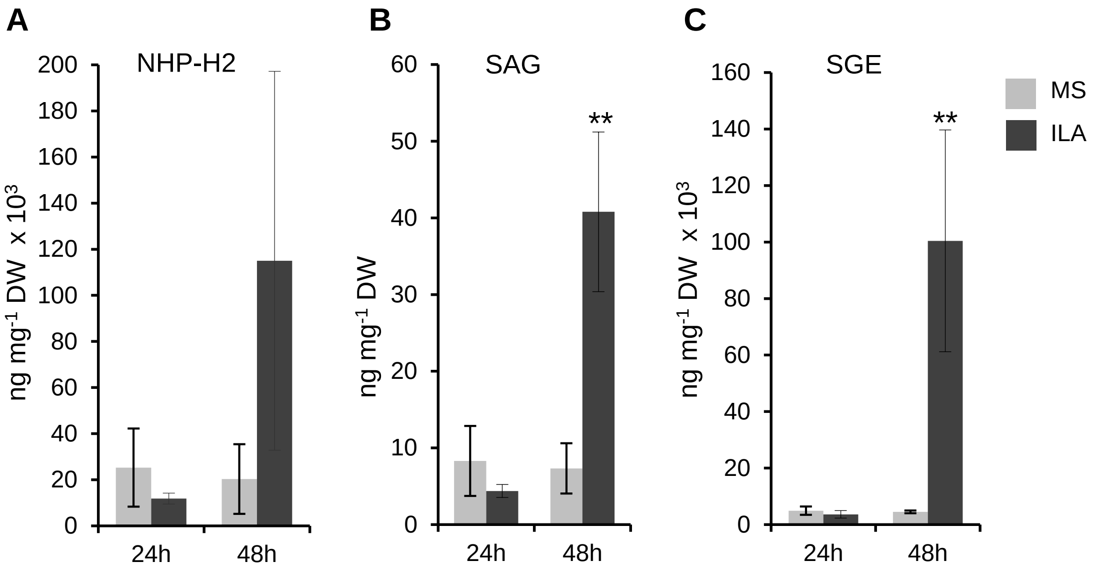


**Supplemental Figure 5**. Exogenous ILA enhances the accumulation of NHP-H2, SAG, and SGE.

**A)** to **(C)** NHP-H2, SAG, and SGE levels of leaves of 12-day-old wild-type seedlings 24 h and 48 h after incubation in ½ MS medium without (grey bar) and with 500 µM ILA (black bars). Bars represent means ± SD; n = 3-4. Asterisks show statistical significance. Differences between treated or untreated plants were analyzed by Welch two sample t-test; * = $p < 0.05$. The p-value for the upregulation of NHP-H2 at 48 h after ILA application was 0.0511.

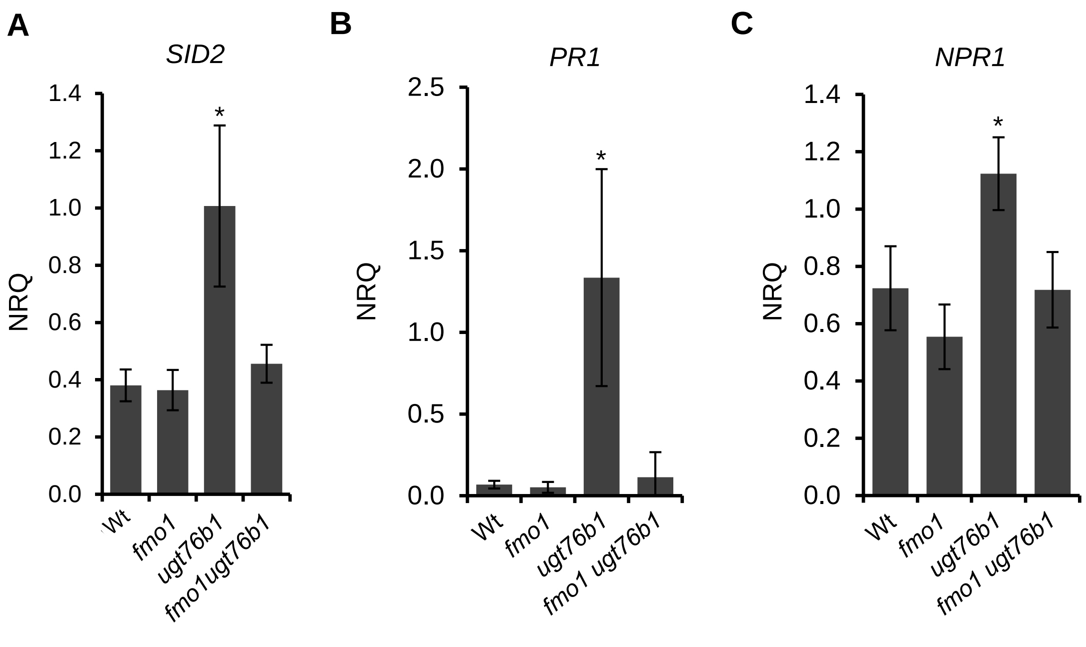


**Supplemental Figure 6**. Introgression of *fmo1* mutation into *ugt76b1* reverses the expression of *SID2* and defense marker genes to wild-type level.

**(A)** to **(C)** RT-qPCR analysis of the transcript abundance of SID2 and the SA-responsive defense marker genes *PR1* and *NPR1* of four-week-old leaves: Gene transcript levels of wild-type, *fmo1*, *ugt76b1,* and *fmo1 ugt76b1* plants grown under short day conditions were analyzed; the normalized relative quantity (NRQ) was determined based on the *UBQ5* and *S16* internal standards; bars show means ± SD, n=4; differences between genotypes were analyzed by Welch two sample t-test; * = $p < 0.05$.
